# Coding Sequence Insertions in Fungal Genomes are Intrinsically Disordered and can Impart Functionally-Important Properties on the Host Protein

**DOI:** 10.1101/2023.04.06.535715

**Authors:** Bernard D. Lemire, Priya Uppuluri

## Abstract

Insertion and deletion mutations (indels) are important mechanisms of generating protein diversity. Indels in coding sequences are under considerable selective pressure to maintain reading frames and to preserve protein function, but once generated, indels provide raw material for the acquisition of new protein properties and functions. We reported recently that coding sequence insertions in the *Candida albicans* NDU1 protein, a mitochondrial protein involved in the assembly of the NADH:ubiquinone oxidoreductase are imperative for respiration, biofilm formation and pathogenesis. NDU1 inserts are specific to CTG-clade fungi, absent in human ortholog and successfully harnessed as drug targets. Here, we present the first comprehensive report investigating indels and clade-defining insertions (CDIs) in fungal proteomes. We investigated 80 ascomycete proteomes encompassing CTG clade species, the Saccharomycetaceae family, the Aspergillaceae family and the Herpotrichiellaceae (black yeasts) family. We identified over 30,000 insertions, 4,000 CDIs and 2,500 clade-defining deletions (CDDs). Insert sizes range from 1 to over 1,000 residues in length, while maximum deletion length is 19 residues. Inserts are strikingly over-represented in protein kinases, and excluded from structural domains and transmembrane segments. Inserts are predicted to be highly disordered. The amino acid compositions of the inserts are highly depleted in hydrophobic residues and enriched in polar residues. An indel in the *Saccharomyces cerevisiae* Sth1 protein, the catalytic subunit of the RSC (Remodel the Structure of Chromatin) complex is predicted to be disordered until it forms a ß-strand upon interaction. This interaction performs a vital role in RSC-mediated transcriptional regulation, thereby expanding protein function.

## IMPORTANCE

We identify thousands of indel mutations that are unique to each of four clades of ascomycete fungi, including several pathogenic species. Indel mutations provide novel insight into the fungal evolutionary processes that may have contributed to the emergence of pathogenicity, metabolic plasticity, and adaptation to extreme environments. Clade-specific mutations offer new avenues into indel-directed drug or protein design to combat fungal pathogenesis or to develop novel bioengineering properties.

## INTRODUCTION

Insertion and deletion mutations (indels) are important evolutionary mechanisms for generating new protein sequences. An indel can substantially change selective pressure on neighboring protein sequences, triggering accelerated evolution that favors adaptation to the presence of the indel (Leushkin et al. 2012). Indels are of great interest because they are known to alter human traits and can cause human diseases (Mills et al. 2006; Bamshad et al. 2011). For example, a single amino acid deletion within the CFTR (cystic fibrosis transmembrane conductance regulator) protein results in cystic fibrosis (Lin et al. 2017). The spike protein of the novel coronavirus SARS-CoV2 contains an insertion ^680^SPRRAR↓SV^687^ that forms a new cleavage site for furin-like proteinases and confers a significant advantage to SARS-CoV2 for entry into human cells (Örd et al., 2020, Shang et al., 2020). Bacterial and protozoan pathogens harbor significant indel content (5-10% in bacteria and up to 25% in protozoa) (Cherkasov, Lee, et al. 2005). The uniqueness of one indel in *Leishmania donovani* elongation factor-1 was used to develop pathogen-specific antibodies (Cherkasov, Nandan, et al. 2005)(Nandan et al. 2007). Thus, there is substantial evidence that indels have a critical role to play in protein evolution as well as in protein interaction networks (Ajawatanawong and Baldauf 2013).

The evolutionary processes that enable pathogenesis and antifungal resistance in yeast are poorly understood. Genome evolution proceeds via many mechanisms, including the acquisition of indels, but very little is known about how genetic variation contributes to virulence, fitness or adaptation in fungi. We previously identified insertion sequences in *Candida albicans* NDU1 that have acquired a function in respiratory growth and virulence in a mouse model of candidiasis (Mamouei et al. 2021). The NDU1 inserts are unique to the CTG-clade fungi. NDU1 is orthologous to NDUFAF6, the NADH:ubiquinone oxidoreductase complex assembly factor 6 (Lemire 2017). Loss of human NDUFAF6 impairs the assembly of mitochondrial complex I (NADH:ubiquinone oxidoreductase) and is associated with Leigh syndrome, a progressive neurodegenerative condition (Baide-Mairena et al. 2019) and with Acadian variant Fanconi Syndrome, a progressive kidney disease (Hartmannová et al. 2016). Expression of the human NDUFAF6 in *C. albicans* did not revert the respiration-defective phenotype of the Ndu1 mutant (Mamouei et al. 2021). This observation suggests that CTG-clade-specific indels in fungal proteins can confer novel virulence-related properties. We recently targeted NDU1 inserts for discovery of a novel antifungal drug niclosamide, with high target specificity, addressing both host toxicity and resistance concerns (Sutar et al. 2022).

The CTG clade fungi are a subdivision of Saccharomycotina (ascomycetes, true yeasts). They acquired a novel genetic code about 170 million years ago (MYa) in which the CTG codon is predominantly translated as a serine instead of a leucine (Massey et al. 2003). Several CTG-clade species are opportunistic pathogens, with *C. albicans, C. auris* and *Candida parapsilosis* being the most common fungal pathogens (Wall et al. 2019). Candidiasis is usually associated with biofilm formation, antifungal resistance and high morbidity. CTG yeasts are also prominent in biotechnology. Species such as *Scheffersomyces stipitis*, *Debaryomyces hansenii and Diutina rugosa* are adept producers of bioethanol, vitamins and other metabolites of medicinal and nutritional value (Riley et al. 2016).

In light of the importance of NDU1 inserts in *C. albicans*, we undertook a study to compare indels in CTG yeasts to closely related fungi. In particular, we were interested in clade-defining inserts (CDIs) and clade-defining deletions (CDDs) that are shared by an evolutionarily-related group of species (Ajawatanawong and Baldauf 2013). The most parsimonious explanation for CDIs and CDDs is that they are the result of mutational events in a common ancestor of the clade that were vertically inherited to the now extant group members (Sharma and Gupta 2019).

Indels identified using protein sequences result from the addition or deletion of a multiple of three nucleotides and preserve the protein reading frame. CDIs are often flanked by conserved blocks of sequences both N- and C-terminally and are mostly restricted to loops on the surface of proteins (Ajawatanawong and Baldauf 2013). The presence of a CDI at a surface location may make it a focal point for acquiring or modifying new intermolecular interactions. In *Saccharomyces cerevisiae*, proteins with indels were found to be involved in more protein-protein interactions (Chan et al. 2007). Indels alter the length of the protein and may allow greater changes in domain orientation, domain packing and protein function than single amino acid substitutions (Emond et al. 2020). Analysis of indels and CDIs in the CTG clade may provide insight into the evolution of pathogenesis, metabolic diversity and adaptation to widely divergent ecological niches.

Indels are abundant in the four clades of ascomycetes we studied. Insertions of five or fewer amino acids account for over 50 % all insertions. Insertions are present in all metabolic pathways and protein families. Proteins oxidative phosphorylation are significantly depleted of insertions in all four clades examined. Conversely, the family of protein kinases is significantly enriched in insertions in all four clades. Insertions are partially excluded from protein structural domains and from transmembrane sequences. Insertion sequences are highly depleted in hydrophobic amino acids, consistent with their predominantly surface exposure on proteins. They are predicted to be largely unstructured but most are flanked by structured regions.

Inserts can modify or expand protein function. One example of this is the *S. cerevisiae* Sth1 protein, which contains two inserts in strategic locations that have key roles in protein-protein interactions. A C-terminal insert in Sth1 provides the recognition site for interaction with the Taf14 transcription factor; loss of this interaction has profound transcriptional effects with significant implications for fitness (Chen et al. 2020). The evolutionary acquisition of inserts can indeed confer novel, functionally important properties on the host protein.

## RESULTS AND DISCUSSION

### Choice of proteomes

We identified evolutionarily conserved indels by comparing four closely related clades of ascomycete fungi. In addition to the CTG-clade yeasts, we chose three other families of Ascomycota with at least 20 available RefSeq proteomes. We analyzed 20 proteomes from the Saccharomycetaceae family; the CTG yeast and Saccharomycetaceae members belong to the subphylum Saccharomycotina, the budding yeasts. In addition, we analyzed 20 proteomes each from the Aspergillaceae and the Herpotrichiellaceae families, which belong to the subphylum Pezizomycotina, or filamentous fungi. Finally, we used the *Schizosaccharomyces pombe* proteome, which belongs to a third subphylum Taphrinomycotina as an outgroup. Protein sequences from 81 yeast strains were obtained and compared (**Figure 1, Supplementary Excel file 1**).

**Figure 1.**
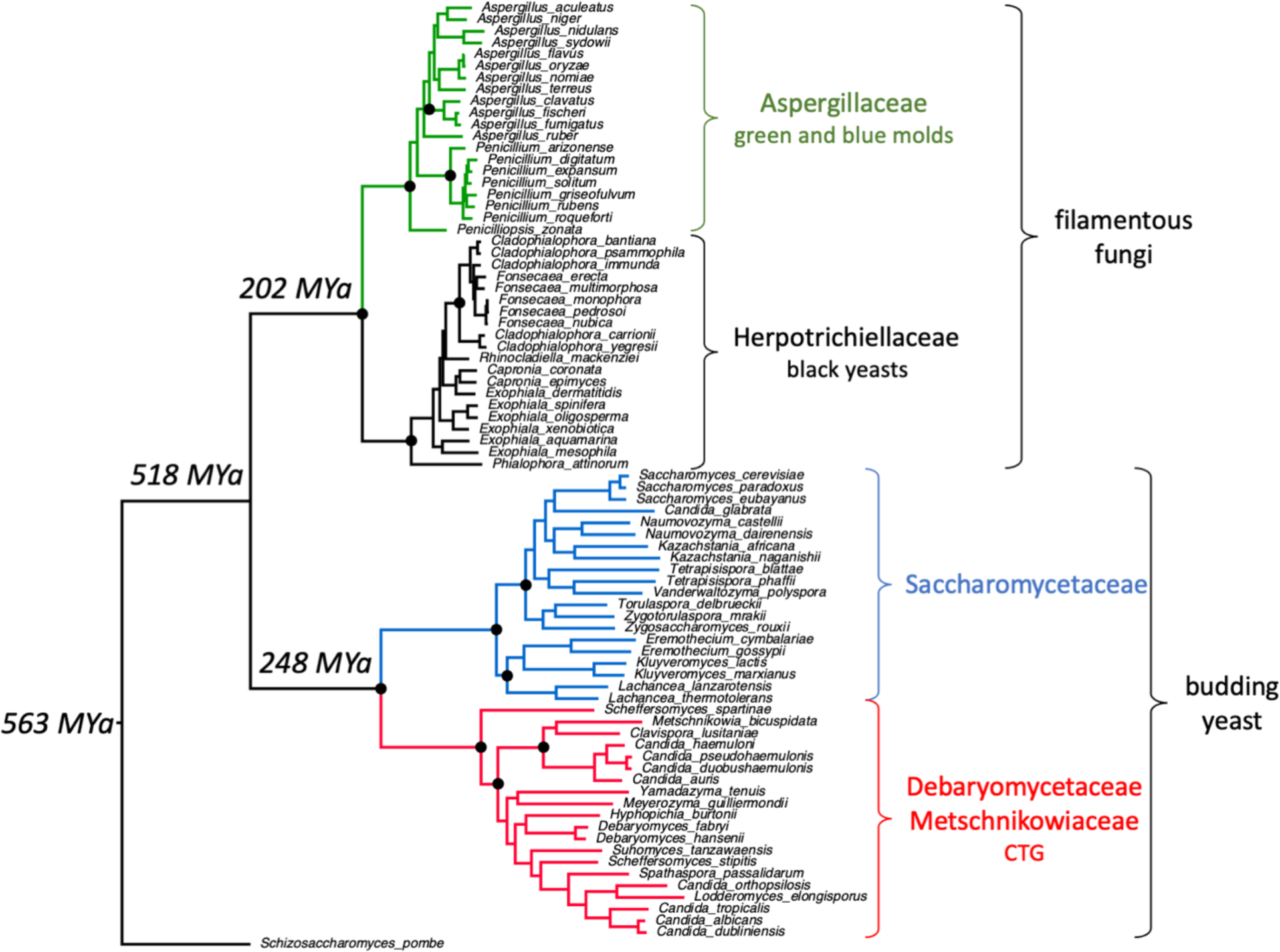
Maximum likelihood phylogeny of 81 ascomycete taxa. 743 multiple sequence alignments each with 81 homologs and containing at least one CDI in the CTG clade were concatenated to produce an alignment of 1,189,784 positions. A maximum likelihood tree was inferred under a Q.yeast+F+I+G4 model using IQ-TREE multicore version 2.1.4-beta (Minh et al. 2020). Selected nodes with 100% bootstrap support are indicated with black circles. All nodes have bootstrap support >60%. The evolutionary times associated with selected nodes were derived from a recent comprehensive investigation of Ascomycota phylogeny (Shen et al. 2020). CUG-Ser1 refers to the CTG clade of yeasts where the tRNACAG^Ser^ can be charged with Leu and result in the translation of CUG codons as leucine (Santos et al. 2011).

The Saccharomycetaceae family includes the model organism *S. cerevisiae*. Family members have proteomes of ∼4,000 to ∼6,000 proteins (**Supplementary Excel file 1**). It also includes *Candida glabrata*, an opportunistic human pathogen that has become a frequent cause of candidemia and is particularly of concern in immunocompromised patients (Gabaldón and Carreté, 2016, Pais et al., 2019).

The Aspergillaceae family are filamentous ascomycetes with greatly expanded capacity for producing secondary metabolites. Many family members are used commercially for food fermentation (*Aspergillus oryzae* for soy sauce, *Penicillium roqueforti* for cheese) or pharmaceuticals (*A. terreus* for lovastatin, *P. rubens* for penicillin) (Houbraken et al. 2020). *A. fumigatus* and *A. flavus* are the most frequently identified *Aspergillus spp.* in human lung infections, causing bronchopulmonary aspergillosis and life-threatening systemic infections in immunocompromised patients (Meena et al. 2021). Aspergillaceae proteomes contain from ∼9,000 to ∼14,000 proteins (**Supplementary Excel file 1**).

The Herpotrichiellaceae family of yeasts belong to the obligatorily melanised black yeasts. Many species are found in extreme or even toxic environments (Teixeira et al. 2017). Some, such as *Fonsecaea pedrosoi* and *Cladophialophora bantiana* are opportunistic pathogens that can cause phaeohyphomycosis (infections by melanised fungi ranging from cutaneous and superficial infections to brain abscesses)(Chowdhary et al. 2015). Their proteomes range in size from ∼9,000 to ∼14,000 proteins (**Supplementary Excel file 1**).

1. *S. pombe* is a member of the fission yeasts and a widely used model organism; in this work, it serves as an outgroup (**Figure 1**). The CTG, Saccharomycetaceae, Aspergillaceae and Herpotrichiellaceae clades had a common ancestor approximately 518 MYa. In addition, the two clades of budding yeast and the two of filamentous fungi diverged from each other at similar times (248 and 202 MYa). Therefore, the accumulation of indels in each clade is on similar timescales.

### Identification and characterization of indels

Our first goal was to investigate indel frequency and length distribution in the four fungal families of Ascomycota. We used *C. albicans SC5314* as our anchor proteome. Each of its 6,030 proteins served as query in a blastp search of the 81 proteomes. A single best hit with an evalue less than 9.9 e^-30^ was retained; blastp hits at this level of significance are highly conserved proteins, often, but not necessarily orthologs (**Supplementary Archive_fasta**). Multiple sequence alignments (MSAs) were generated for each query protein and its blastp hits meeting the evalue threshold (**Supplementary Archive_MSAs**). Of the 6,030 MSAs generated, 1,532 MSAs (25.4% of *C. albicans* proteins) contained a full set of 81 homologous protein sequences and only these MSAs were analyzed. These criteria are stringent and remove many important proteins from our analysis. Shorter proteins require greater sequence identity to attain the evalue threshold necessary for inclusion. The shortest *C. albicans* protein included in the set of 1,532 MSAs is 148 amino acids in length. Proteins belonging to missing pathways will be excluded. One example is NDU1; Saccharomycetaceae family members have lost complex I and do not have homologs of NDU1 (Pagliarini et al. 2008). The MSA for NDU1 has only 34 homologs and is not included in the indel analysis. These strict criteria were deliberately chosen so that 20 sequences are present from each of the four taxonomic families and that these proteins are unquestionably homologous.

A python program was used to identify clade-or strain-specific inserts present in the MSAs. N-and C-terminal indels were excluded. Mechanistically, the generation of N- and C-terminal indels differ from the generation of internal indels. For example, N-terminal indels may be generated through the use of alternative translation start codons, while C-terminal indels may be generated by frameshift mutations generating alternative stop codons; neither of these mechanisms will generate an internal indel.

Inserts are abundant in ascomycete proteomes. Over 26,000 insertions were found in the 1,532 MSAs (**Table 1, Supplementary Excel file 5**). Most proteins have insertions. For example, the CTG-specific inserts are found in 1,308 MSAs or 85.4% of the 1,532 proteins analyzed. Of the 1,532 MSAs, only 15 do not have any inserts in any of the four clades. There are fewer inserts in the Aspergillaceae and the Herpotrichiellaceae clades (∼60%) than in the CTG and Saccharomycetaceae clades (**Table 1**). The total number of inserts in each clade correlates with the rate of evolutionary change. The number of inserts per unit branch length to the last common ancestor of all 4 clades (518 MYa in **Figure 1**) for the CTG, Saccharomycetaceae, Aspergillaceae and Herpotrichiellaceae clades, respectively are 0.50, 0.53, 0.51 and 0.48. The longer branch lengths in Saccharomycotina (CTG and Saccharomycetaceae) compared to Pezizomycotina (Aspergillaceae and Herpotrichiellaceae) are indicative of higher Saccharomycotina evolutionary rates (Shen et al. 2020).

**Table 1.** Number of inserts in each of the four taxonomic families. All inserts except those at the N-or C-termini were counted. The total number of MSAs analyzed is 1,532 and the number of MSAs in which all insertions are found is indicated. Only MSAs with inserts are included in the calculation of inserts per MSA. Sacc (Saccharomycetaceae) clade; Asp (Aspergillaceae) clade; Herpo (Herpotrichiellaceae) clade.

| **Taxonomic family** | **CTG** | **Sacc** | **Asp** | **Herpo** |
| --- | --- | --- | --- | --- |
| no. of inserts total | 7,418 | 8,481 | 5,301 | 5,031 |
| insert length 1 | 1,969 | 2,426 | 1,167 | 1,259 |
| insert length 1-5 | 4,241 | 5,225 | 2,712 | 2,985 |
| insert length 6-10 | 1,006 | 1,241 | 743 | 756 |
| insert length 11-20 | 858 | 926 | 815 | 613 |
| insert length 21-50 | 715 | 734 | 485 | 431 |
| insert length >50 | 598 | 355 | 486 | 245 |
| insert length >100 | 288 | 91 | 268 | 125 |
| insert length 101-200 | 165 | 77 | 143 | 62 |
| insert length 201-300 | 51 | 9 | 60 | 26 |
| insert length 301-400 | 24 | 5 | 27 | 16 |
| insert length 401-500 | 11 | 0 | 15 | 6 |
| insert length 501-600 | 12 | 0 | 10 | 2 |
| insert length 601-700 | 7 | 0 | 4 | 1 |
| insert length 701-800 | 4 | 0 | 2 | 3 |
| insert length 801-900 | 3 | 0 | 4 | 4 |
| insert length 901-1000 | 1 | 0 | 1 | 1 |
| >1001 | 10 | 0 | 2 | 4 |
| no. of MSAs with inserts | 1,308 | 1,349 | 1,288 | 1,266 |
| inserts per MSA | 5.67 | 6.29 | 4.12 | 3.97 |

Single amino acid inserts comprise 26.0% of all inserts, and inserts of 5 or fewer residues in length 57.8% (**Table 1, Figure 2A**). These short inserts are the most highly abundant for all clades. Although the frequency of longer inserts decreases rapidly, it is noteworthy that inserts of hundreds of amino acids can be found. The Saccharomycetaceae clade has the greatest total number of inserts, but longer inserts are fewer in this clade, with none greater than 400 residues in length (**Table 1**). In contrast, the CTG, Aspergillaceae and Herpotrichiellaceae clades have 48, 38 and 21 inserts >400 residues, respectively.

**Figure 2A.**
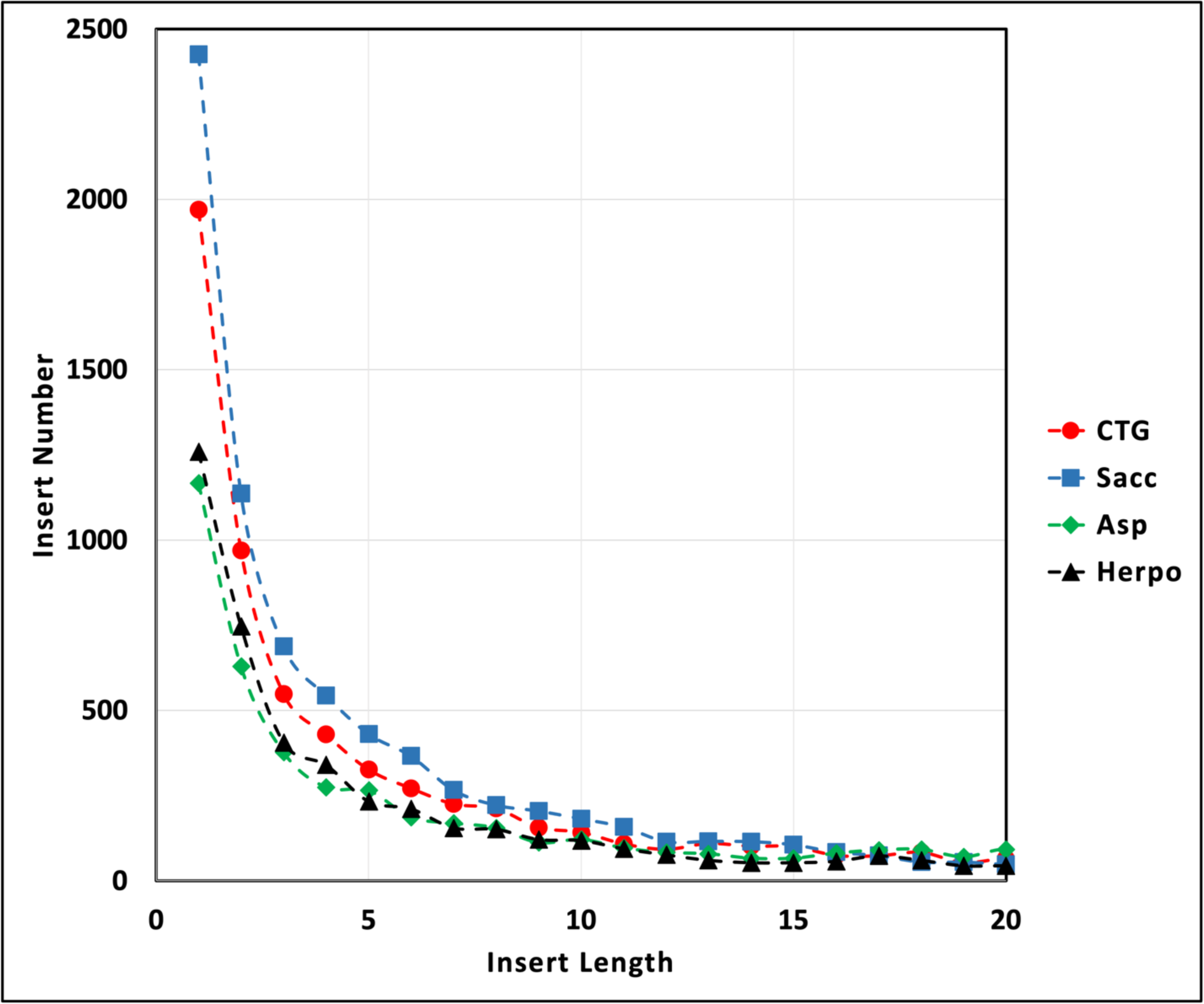
Length distribution of short inserts. The numbers of inserts for insert lengths from 1 to 20 are plotted for each clade. Red circles, CTG clade; blue squares, Saccharomycetaceae (Sacc) clade; green diamonds, Aspergillaceae (Asp) clade; black triangles, Herpotrichiellaceae (Herpo) clade.

### Clade-defining insertions and deletions

We classified insertion events as algorithm-related inserts, singleton inserts and CDIs. For algorithm-related inserts, the length or even the presence of an insert is dependent on algorithm parameters, such as the scoring matrix and gap-insertion penalties (**Supplementary Figure 1A**). These inserts generally do not have well-aligned blocks of residues upstream and downstream of the insert. Algorithm-related inserts are less likely to be biologically relevant and we focused our analysis on CDIs.

We defined CDIs as having at least one flanking column with a minimum of 59 amino acids on each side of the insert; this ensures proteins from all 4 clades show conservation in the flanking positions (**Supplementary Figure 1A**). We also chose to focus our analysis on CDIs with at least one insert position having a residue in 5 or more strains of the same clade. CDIs shared by multiple strains are more evolutionarily informative, having arisen in a common ancestor and being vertically inherited. A subclass of CDIs are the singleton inserts; they are found in a single taxon and may have arisen after speciation (**Supplementary Figure 1A**).

Over 4,000 CDIs were found in the 1,532 MSAs analyzed (**Table 2, Supplementary Excel file 4**). The CTG and Saccharomycetaceae clades have ∼3 to ∼5 times more CDIs than do the other two clades. They have more proteins with CDIs and a greater number of CDIs per protein. CDIs account for 22.0%, 20.6%, 6.8% and 11.0% of all inserts in the CTG, Saccharomycetaceae, Aspergillaceae and Herpotrichiellaceae clades, respectively. The differences in numbers of CDIs between clades cannot be explained by differing evolutionary rates, as for the total numbers of inserts. The larger numbers of CDIs in the CTG and Saccharomycetaceae may arise from different repair rates of the original mutational events that gave rise to the CDIs. Shen *et al.,* have reported that, on average, Saccharomycotina (CTG and Saccharomycetaceae) have fewer DNA repair genes (average 41) than Pezizomycotina (Aspergillaceae and Herpotrichiellaceae; average 54) (Shen et al. 2020).

**Table 2.** Number of CDIs and CDDs in each of the four taxonomic families. CDIs must have flanking positions with 59 or more residues and at least one position in the insert with 5 residues. CDDs have no amino acids in the clade of interest, at least five amino acids in each of the other three clades and at least 40 residues in the flanking positions. CDIs and CDDs at the N-or C-termini were not counted. The total number of MSAs analyzed is 1,532 and the number of MSAs in which the CDIs and CDDs are found is indicated. Only MSAs with CDIs or CDDs are included in the calculation of CDIs per MSA or CDDs per MSA. Sacc (Saccharomycetaceae) clade; Asp (Aspergillaceae) clade; Herpo (Herpotrichiellaceae) clade.

| **Taxonomic family** | **CTG** | **Sacc** | **Asp** | **Herpo** |
| --- | --- | --- | --- | --- |
| CDIs total | 1,630 | 1,743 | 360 | 555 |
| No. of proteins with CDIs | 743 | 783 | 285 | 368 |
| CDIs per protein | 2.19 | 2.23 | 1.26 | 1.51 |
| CDI length 1-10 | 1,294 | 1,372 | 268 | 453 |
| CDI length 11-20 | 166 | 180 | 34 | 51 |
| CDI length 21-30 | 64 | 72 | 13 | 21 |
| CDI length 31-40 | 32 | 43 | 7 | 7 |
| CDI length 41-50 | 16 | 22 | 5 | 11 |
| CDI length 51-100 | 39 | 43 | 6 | 5 |
| CDI length >101 | 19 | 11 | 27 | 7 |
| CDDs total | 633 | 687 | 596 | 603 |
| No. of proteins with CDDs | 434 | 439 | 429 | 423 |
| CDDs per protein | 1.46 | 1.56 | 1.39 | 1.43 |
| CDDs length 1-5 | 600 | 648 | 573 | 578 |
| CDDs length 6-10 | 31 | 28 | 21 | 24 |
| CDDs length >11 | 2 | 11 | 2 | 1 |

The majority of CDIs are short. CDIs of one amino acid comprise approximately one third of all CDIs (**Table 2, Figure 2B**). CDIs of 10 or fewer residues account for 74-82% of all CDIs in each clade. Longer CDIs are presumably more disruptive of the host protein’s structure and/or function and subject to selective elimination.

**Figure 2B.**
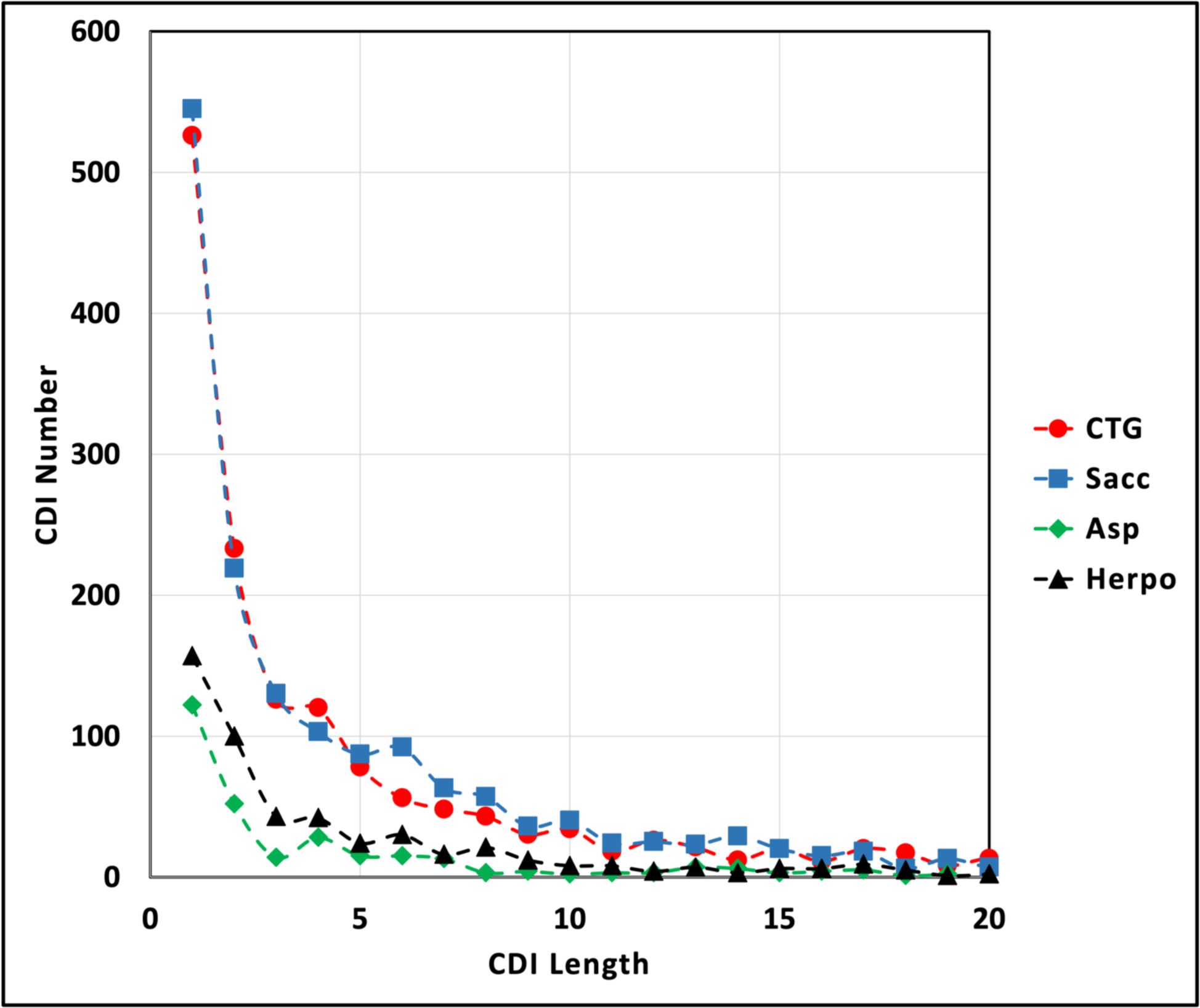
Length distribution of short CDIs. The numbers of CDIs up to a length of 20 were plotted for each clade. Red circles, CTG clade; blue squares, Sacc (Saccharomycetaceae) clade; green diamonds, Asp (Aspergillaceae) clade; black triangles, Herpo (Herpotrichiellaceae) clade.

We defined CDDs as having no amino acids in the clade under investigation, at least 5 aligned amino acids in each of the three other clades and flanking columns of 40 or more aligned residues on both sides of the deletion (**Supplementary Figure 1B**). Algorithm-related deletions can also be created by the way the algorithm inserts gaps into poorly aligning regions of MSAs. Over 2,500 CDDs were found in the 1,532 MSAs analyzed (**Table 2, Supplementary excel file 4**). CDDs are much more restricted in length than CDIs, reaching a maximum length of 19 (**Supplementary Figure 2A**). Short deletions of fewer than five amino acids comprise an even greater majority than short CDIs. The number of CDDs and the number of CDDs per protein are roughly similar in all four clades. In contrast, the number of CDIs is considerable greater in the CTG and Saccharomycetaceae clades than in the Aspergillaceae and Herpotrichiellaceae clades, where CDDs outnumber CDIs. These observations suggest that insertions and deletions may arise through mechanistically different processes.

### CDI generation and retention

CDIs are prominent features that help define clades. Each of the four distinct clades has its own distinct set of CDIs that most likely originated in a common ancestor of the clade members after the split from sister clades had occurred. The number of shared CDIs in a clade provides an estimate of the evolutionary rate of CDI loss. The rates of CDI loss range from 0.7-3.2 CDIs/MY. These rates differ as much between strains in the same clade as between clades (**Supplementary Table 1, Supplementary Figure 2B**)

We also determined the numbers of CDIs shared by the budding yeast in the Saccharomycotina clade and by the filamentous fungi in the Pezizomycotina clade. CDIs shared by the budding yeasts were generated at ∼3.2 inserts/MY (853 CDIs/270 MY); filamentous fungal inserts were generated at ∼2.5 inserts/MY (784 CDIs/316 MY). The rate of CDI loss (0.7-3.2 CDIs/MY) is comparable to the rate of generation (2.5-3.2 inserts/MY). The most parsimonious explanation for these CDIs is that the insertion event occurred after the last common ancestor of all the strains (518 MYa), but before the split between the Saccharomycotina and the Pezizomycotina clades (248 MYa for the budding yeast and 202 MYa for the filamentous fungi) (**Supplementary Figure 2B**).

### *C. albicans* inserts are partially excluded from structural domains and strongly excluded from transmembrane domains

Domains are distinct functional or structural units of proteins and usually fold independently of other parts of the protein. We investigated the locations of inserts and domains in *C. albicans* proteins to determine whether insert frequency is affected by the presence of domains. Within the set of 1,532 MSAs, we identified the inserts present in the *C. albicans* homolog. 3,069 inserts were found in 1,015 MSAs (3.0 inserts per *C. albicans* protein). Over 30 proteins harbor 10-21 inserts per protein.

The 1,015 *C. albicans* proteins with inserts were analyzed for Pfam (Pfam 35.0) domains (Mistry et al. 2021). Of these, 984 proteins (96.9%) contained 1,884 Pfam domains with an evalue less than 9.99e-06 (1.9 domains/protein). Inserts and Pfam domains were plotted (**Figure 2C**). Of the 3,069 *C. albicans* inserts, 1,331 (43.4%) occur within 800 Pfam domains, yet these domains account for 51.9% of the protein lengths. These data indicate that inserts are somewhat excluded from being in domains.

**Figure 2C.**
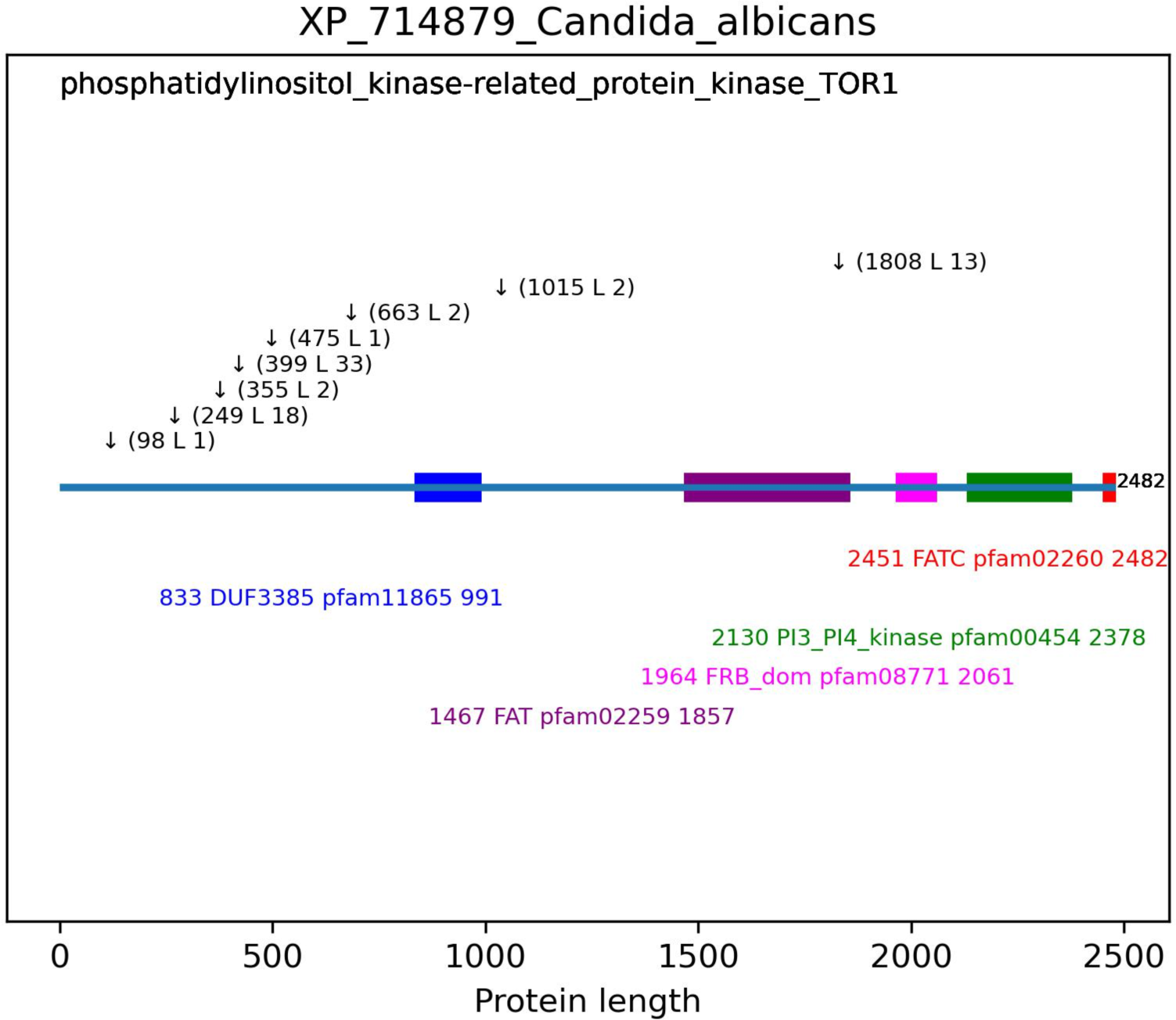
Schematic representation of domains and inserts. *C. albicans* protein sequences with inserts were submitted to Conserved Domain Database for domain analysis using the Pfam database (Lu et al. 2020). The length of the protein is indicated at the end of the line depicting the protein. Domains are depicted as colored rectangles with matching colored text below indicating the domain start site, the domain name, the Pfam domain ID and the domain end site. Inserts are indicated by the downward arrows followed by the insert start site and the length of the insert. In the top left corner is the protein description from the fasta file.

Transmembrane segments (TMs) are important structural domains present in a hydrophobic environment. There may be increased selective pressure to remove inserts from TMs because they have specific hydrophobicity and secondary structure requirements. In addition, TMs may interact with each other or with TMs of other proteins and inserts may result in interacting surfaces facing in incorrect directions. We analyzed 1,532 *C. albicans* proteins in the MSA pool for TMs using DeepTMHMM, an algorithm that uses deep learning to predict TMs (Hallgren et al. 2022). We found 2,368 TMs in 315 proteins (20.6% of the total). Of the 315 membrane proteins, 219 contain 593 inserts (69.5% of membrane proteins have inserts). When we analyzed the positions of inserts with respect to TMs, we found 27 inserts were located in the predicted TMs. The fraction of protein length occupied by 2,368 TMs is 22.06%, yet inserts are found in only 4.55% of TMs, indicating that inserts are strongly excluded from TMs. As presented below, inserts are largely hydrophilic and unstructured, both properties that are incompatible with the formation of TMs.

Proteins with inserts are longer than the average protein in *C. albicans* (**Figure 2D**). The 6,030 proteins of the proteome have an average length of 494 amino acids/protein. Insert-containing proteins have an average of 707 amino acids per protein and range in length from 148 to 3,286 amino acids.

**Figure 2D.**
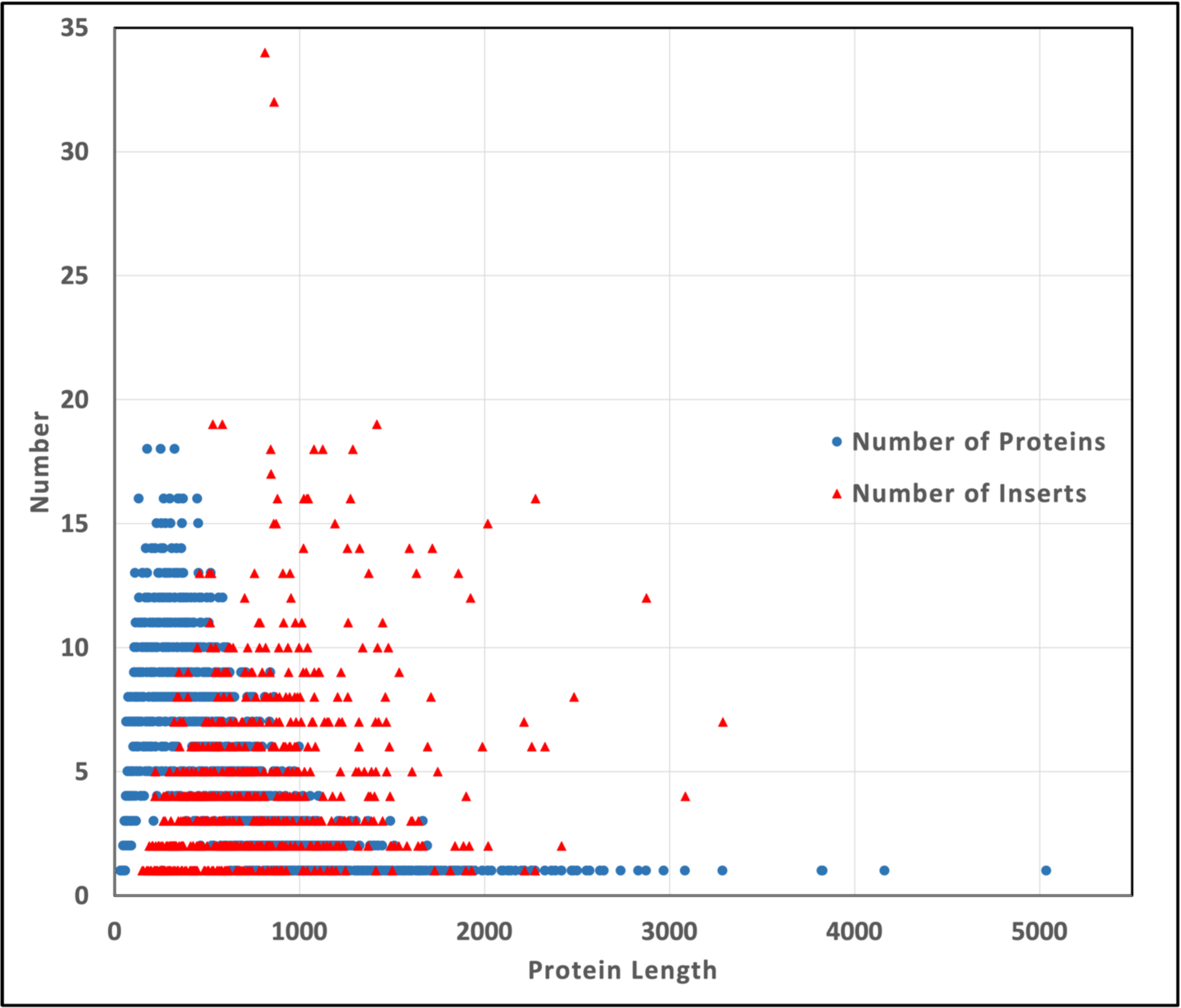
The numbers of proteins and inserts as a function of protein length. The 6,030 proteins of the *C. albicans* proteome are plotted by protein length (blue circles). The distribution of the 3,069 inserts identified in 1,015 *C. albicans* proteins is plotted against protein length (red triangles).

Few structures of *C. albicans* proteins have been determined. One that has is the C-terminal domain (residues 702-1107) of the *C. albicans* trehalose synthase regulatory protein TPS3 (XP_019330821, UniProt accession A0A1D8PIS4, PDB:5HUS). It contains a Pfam02358, trehalose_PPase domain and four inserts (**Figure 2E**). The longest insert (731-743) is only partly modeled in the structure, suggesting it is mobile and unstructured. The two inserts within the Pfam domain are short (one and two amino acids, respectively) and are present at the ends of a helix in loops, where they have little effect on the overall domain structure. As with the majority of inserts, the TPS3 inserts are exposed on the protein surface of the protein.

**Figure 2E.**
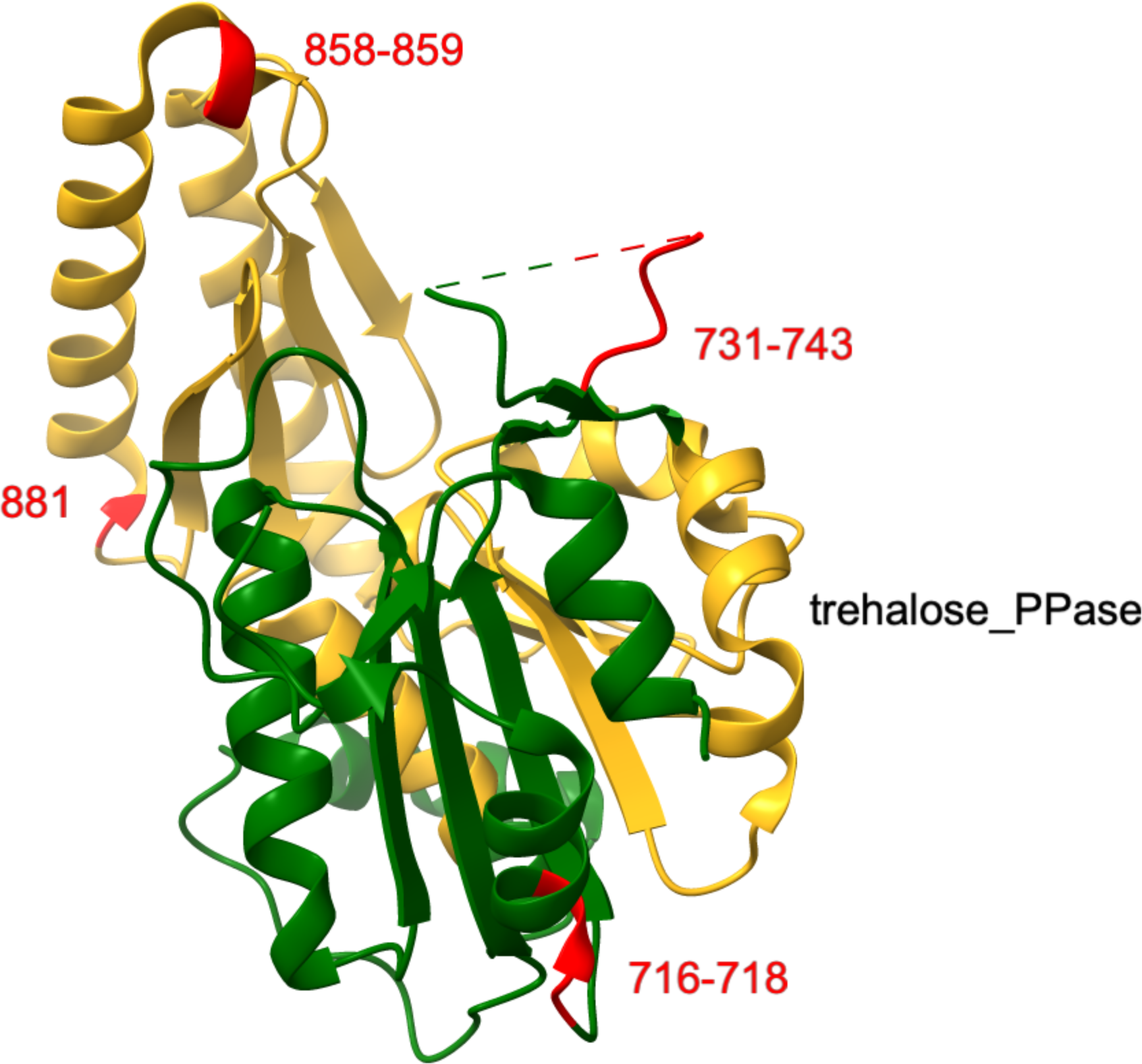
The structure of the *C. albicans* trehalose 6-phosphate synthase/phosphatase complex subunit. The C-terminal domain (residues 702-1107) of the *C. albicans* protein XP_019330821 (UniProt accession A0A1D8PIS4) is shown in green in the structure PDB:5HUS (unpublished). The Pfam domain pfam02358, trehalose_PPase (residues 754-911) is colored gold. The positions of four inserts are indicated and corresponding residues are colored red.

### Orthology analysis of insert-containing proteins

What types of proteins have inserts? To answer this question, one strain from each clade was chosen for orthology analysis. KofamKOALA assigns protein sequences to KEGG Ortholog families (KOfam) by using profile hidden Markov models and link protein sequences to KEGG pathways (Aramaki et al. 2020). Proteins with inserts from *C. albicans*, *S. cerevisiae*, *A. oryzae* and *C. coronata* were identified were submitted for orthology assignment (**Supplementary Table 2, Supplementary Excel file 3**). For *C. albicans*, *S. cerevisiae*, *A. oryzae* and *C. coronata*, 84.3%, 85.2%, 88.8% and 90.2% of submitted sequences were successfully assigned to a KEGG ortholog family, respectively, and 57.3%, 58.4%, 61.1% and 61.1% assigned to five functional classifications and specific pathways (**Table 3, Supplementary Excel file 2**).

**Table 3.**
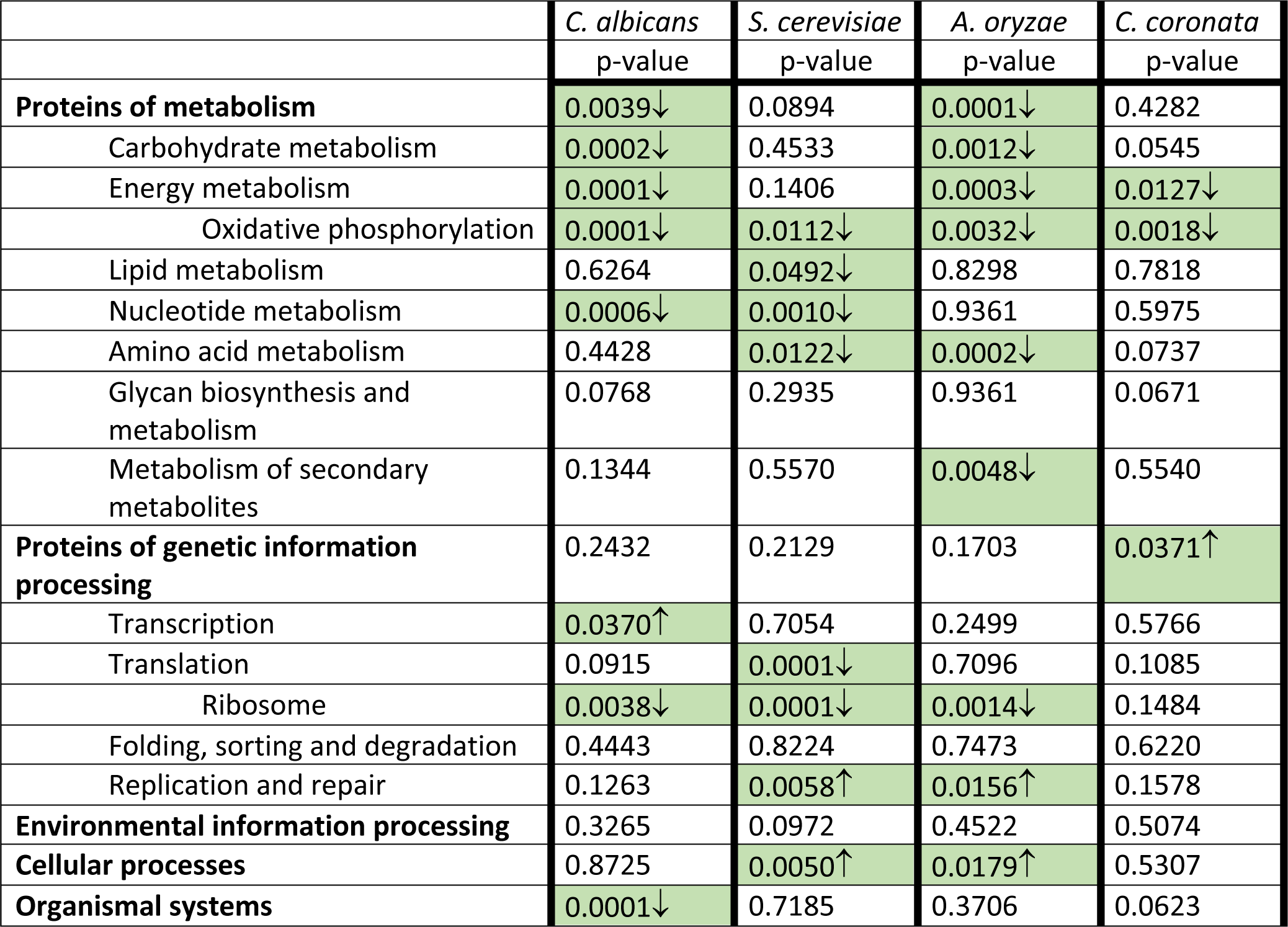
Pathway analysis comparing insert-containing proteins to the entire pool of aligned sequences. Insert-containing proteins and the pool of 1,532 proteins analyzed were submitted to KEGG for orthology assignment. The proteins were grouped into five global pathway categories (bold listings). The numbers of proteins included in each category or pathway for insert-containing proteins and for proteins in the pool were compared using Chi-square analysis. The null hypothesis is that membership in a pathway and the presence of inserts are independent variables. pvalues < 0.0001 are recorded as 0.0001. pvalues < 0.05 indicate the null hypothesis is rejected and are highlighted in green. Up and down arrows in highlighted cells indicate comparisons where more or fewer insert-containing proteins belong to the category than expected, respectively.

|  | *C. albicans* | *S. cerevisiae* | *A. oryzae* | *C. coronata* |
| --- | --- | --- | --- | --- |
|  | p-value | p-value | p-value | p-value |
| **Proteins of metabolism** | 0.0039↓ | 0.0894 | 0.0001↓ | 0.4282 |
| Carbohydrate metabolism | 0.0002↓ | 0.4533 | 0.0012↓ | 0.0545 |
| Energy metabolism | 0.0001↓ | 0.1406 | 0.0003↓ | 0.0127↓ |
| Oxidative phosphorylation | 0.0001↓ | 0.0112↓ | 0.0032↓ | 0.0018↓ |
| Lipid metabolism | 0.6264 | 0.0492↓ | 0.8298 | 0.7818 |
| Nucleotide metabolism | 0.0006↓ | 0.0010↓ | 0.9361 | 0.5975 |
| Amino acid metabolism | 0.4428 | 0.0122↓ | 0.0002↓ | 0.0737 |
| Glycan biosynthesis and metabolism | 0.0768 | 0.2935 | 0.9361 | 0.0671 |
| Metabolism of secondary metabolites | 0.1344 | 0.5570 | 0.0048↓ | 0.5540 |
| **Proteins of genetic information processing** | 0.2432 | 0.2129 | 0.1703 | 0.0371↑ |
| Transcription | 0.0370↑ | 0.7054 | 0.2499 | 0.5766 |
| Translation | 0.0915 | 0.0001↓ | 0.7096 | 0.1085 |
| Ribosome | 0.0038↓ | 0.0001↓ | 0.0014↓ | 0.1484 |
| Folding, sorting and degradation | 0.4443 | 0.8224 | 0.7473 | 0.6220 |
| Replication and repair | 0.1263 | 0.0058↑ | 0.0156↑ | 0.1578 |
| **Environmental information processing** | 0.3265 | 0.0972 | 0.4522 | 0.5074 |
| **Cellular processes** | 0.8725 | 0.0050↑ | 0.0179↑ | 0.5307 |
| **Organismal systems** | 0.0001↓ | 0.7185 | 0.3706 | 0.0623 |

Although our investigations into inserts originated from our observations on the *C. albicans* NDU1 (Mamouei et al. 2021), the data in **Table 3** indicate that inserts are not unique to *C. albicans*. Oxidative phosphorylation is the only pathway that shows a statistically significant depletion of insert-containing proteins in all four strains (**Table 3, Supplementary Excel file 2**). Proteins of oxidative phosphorylation may have experienced fewer insertion events or have a lower retention rate or both. One functional category, organismal systems and one individual pathway, transcription, show *C. albicans*-specific depletion of inserts. Organismal systems includes proteins associated with the immune, endocrine, circulatory, digestive, excretory, sensory and nervous systems. Examples of strain-specific enrichment or depletion of inserts in pathways can also be found in each of the other three strains, again indicating that *C. albicans* inserts are not unique.

Where a pathway shows a statistically significant increase or decrease in insert-containing proteins in more than one strain, the pattern is always consistent across the strains involved. All strains will either show a decrease or an increase in insert-containing proteins. These observations suggest that the presence of inserts in proteins may be determined partly by the nature of the pathways or the protein complexes they belong to.

KEGG also assigns proteins to protein families (**Table 4, Supplementary Excel file 2**). Only one family, the protein kinases, is significantly enriched in insert-containing proteins in all four strains. In *C. albicans*, *S. cerevisiae*, *A. oryzae* and *C. coronata*, 82.4%, 87.8%, 71.4% and 71.2% of protein kinases have inserts, respectively. This family has either experienced more insertion events or had a higher insert retention rate or both over the last ∼200 MY. Indels are a common mechanism of kinase activation in cancer—a feature exploited clinically by targeted therapy with kinase inhibitors (Sehn 2015). In fact, indels account for 18% of the sequence difference between mice and human kinomes (Caenepeel et al. 2004). The presence of more inserts in kinases suggests that new insertion events did not abolish protein function. Two protein families, glycosyltransferases and proteasomal proteins show *C. albicans*-specific enrichment and depletion of inserts, respectively.

**Table 4.**
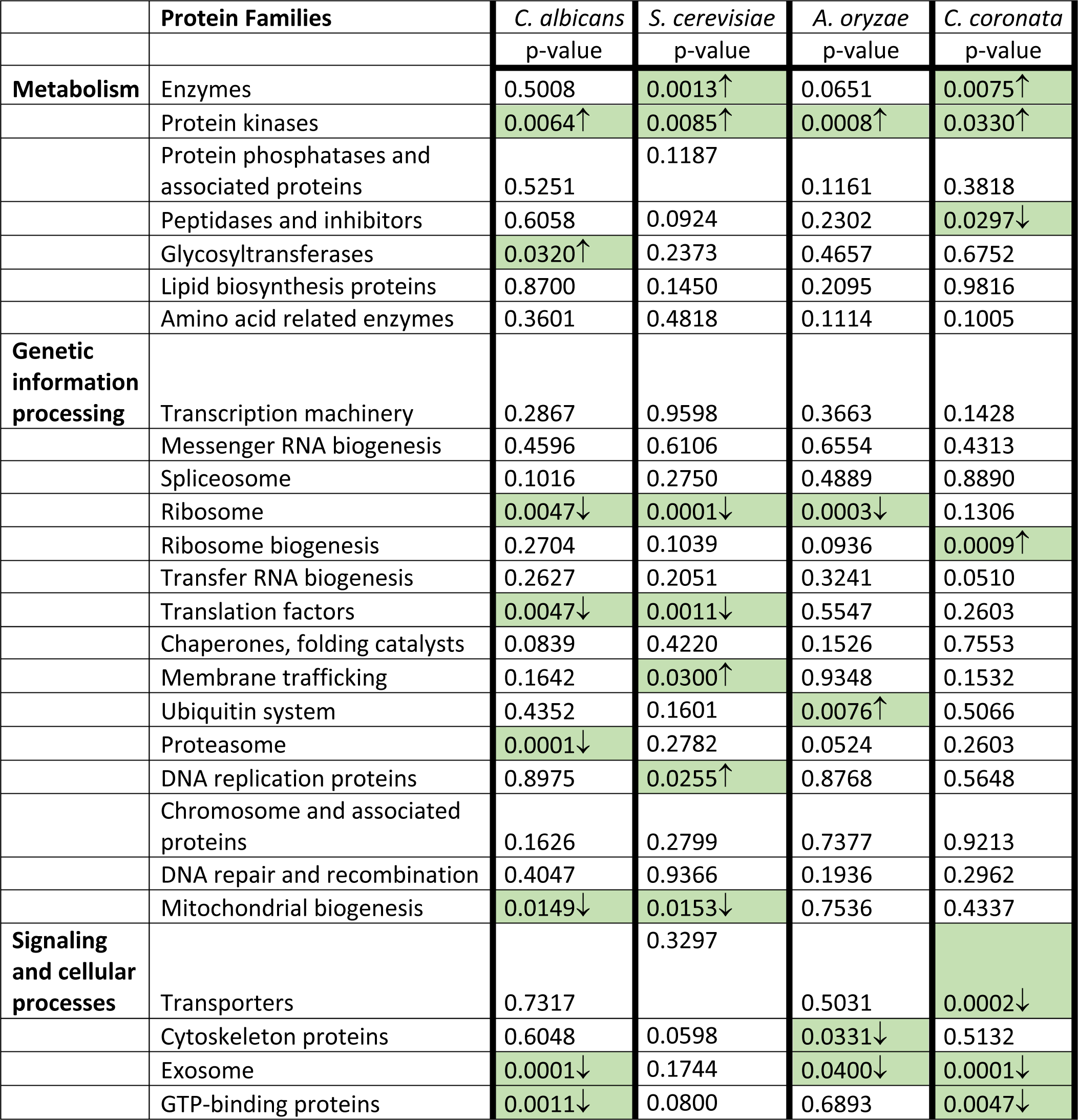
Protein family analysis comparing insert-containing proteins to the entire pool of aligned sequences. Insert-containing proteins and the pool of 1,532 proteins were submitted to KEGG for orthology assignment. The proteins were grouped into three functional categories of proteins (bold listings). The numbers of proteins included in each protein family for insert-containing proteins and for proteins in the pool were compared using Chi-square analysis. The null hypothesis is that membership in a pathway and the presence of inserts are independent variables. pvalues < 0.0001 are recorded as 0.0001. pvalues < 0.05 indicate the null hypothesis is rejected and are highlighted in green. Up and down arrows in highlighted cells indicate comparisons where more or fewer insert-containing proteins belong to the category than expected, respectively.

### *C. albicans* and *S. cerevisiae* essential proteins have fewer inserts

Essential proteins in *S. cerevisiae* were found to have a higher indel frequency than non-essential proteins and the indel-containing proteins were involved in more protein-protein interactions (Chan et al. 2007). We investigated the frequency of inserts in essential genes.

The Saccharomyces Genome Database annotates 1,313 genes as essential (Engel et al. 2021). Of these, 1,239 could be mapped to RefSeq accessions. Of the 968 insert-containing *S. cerevisiae* proteins, 322 are encoded by essential genes (33.3%). In the pool of 1,532 MSAs analyzed, 1,355 unique *S. cerevisiae* proteins are present, with 467 (34.5%) of these being essential. In *S. cerevisiae*, the presence of inserts is independent of the gene being essential, χ^2^ (1, *N=1355)* 2.2, p=.14. These results are not in agreement with the results of Chan *etal*., 2017. One notable methodological difference may account for the different results. While our MSAs include 81 ascomycete proteins, the *S. cerevisiae* MSAs in the previous study include 15 additional eukaryotic species that are taxonomically much more diverse and include *Arabidopsis thaliana*, *Caenorhabditis elegans* and *Trypanosoma cruzi* (Chan et al. 2007).

A list of 1,660 genes annotated with an inviable phenotype was obtained from the Candida Genome Database (Skrzypek et al. 2018). Of these, 1,654 could be mapped to RefSeq accessions. Of the 1,015 insert-containing *C. albicans* proteins, 385 are encoded by essential genes (37.9%). Of the 1,532 MSAs analyzed, 648 (42.3%) are essential genes. There are significantly fewer inserts in essential genes than expected, χ^2^ (1, *N=1532)* = 23.5, p=1.3e^-6^. The exclusion of inserts from *C. albicans* essential proteins is not seen in *S. cerevisiae*. Unlike *S. cerevisiae, C. albicans* is a heterozygous diploid yeast that relies on oxidative metabolism (Jones et al. 2004) and this different metabolism may account for a significant proportion of the additional number of essential proteins in *C. albicans* (1,660) compared to *S. cerevisiae* (1,313).

### *C. albicans* and *S. cerevisiae* insert-containing proteins do not have more protein-protein interactions

Does the presence of inserts in essential proteins affect the number of protein-protein interactions? We used the STRING database of known and predicted protein-protein interactions to determine the number of nodes (interacting proteins) and edges in the physical subnetwork, where the edges are the evidence that indicates a protein is part of a physical complex (Szklarczyk et al. 2021). The essential proteins of *C. albicans* and *S. cerevisiae* with and without inserts were submitted to STRING for analysis; the interaction results were limited to those of highest confidence (score>0.9). A two sample t-test was performed to compare the number of nodes of *C. albicans* essential proteins with or without inserts. There was not a significant difference in number of nodes between proteins with inserts (N = 385, M = 7.27, SD = 4.18) and proteins without inserts (N = 262, M = 7.69, SD = 4.12); t(22) = −1.26, p =.21. Similarly, when a two sample t-test was performed to compare the number of edges, there was not a significant difference between proteins with inserts (M = 24.95, SD = 21.74) and proteins without inserts (M = 25.07, SD = 21.70); t(22) = −0.07, p =.95. When a t-test to compare the number of nodes in *S. cerevisiae* essential proteins was performed, no significant difference was found between proteins with inserts (N = 322, M = 6.87, SD = 4.20) and proteins without inserts (N = 145, M = 7.12, SD = 4.41); t(22) = −0.57, p =.57. For comparing the number of edges in *S. cerevisiae* essential proteins, the t-test was not significant for proteins with inserts (M = 24.39, SD = 22.22) and proteins without inserts (M = 25.63, SD = 22.74); t(22) = −0.55, p =.58. Inserts do not increase the number of protein-protein interactions in *C. albicans* or in *S. cerevisiae.* Our results do not concur with those of Chan *et al.,* 2007, but again, significant methodological differences may account for the divergence. That inserts do not significantly increase the numbers of protein-protein interactions does not mean that no inserts participate in protein-protein interactions. An example of inserts playing key roles in protein oligomerization is *S. cerevisiae* Sth1, discussed below.

### *S. cerevisiae* insert-containing proteins do not form more complexes

Complex Portal is a database of macromolecular complexes (Meldal et al. 2021). It comprises 589 complexes with over 1,900 proteins. *S. cerevisiae* has 322 essential proteins with inserts, of which 157 (48.8%) are found in Complex Portal. An additional 145 essential proteins do not have inserts and 59 of these are in Complex Portal (40.7%). Chi-square analysis indicates that the presence of inserts and participation in a complex are independent variables, χ^2^ (1, *N=467*) = 2.6, p=.11). Similarly, when all insert-containing proteins are analyzed for their presence in complexes, they are not over-represented in complexes. *S. cerevisiae* has 968 insert-containing proteins, with 313 (32.3%) in Complex Portal. Of the 387 *S. cerevisiae* proteins without inserts, 119 (30.7%) are present in complexes, χ^2^ (1, *N=1355*) = 0.32, p=.57). We conclude that inserts do not promote complex formation.

### Insertion sequences are disordered

Do inserts fold into stable structures? Does insert structure depend on insert length and on the structure of the insertion site? Very few structures for *C. albicans* proteins have been determined. When structures are available, insert sequences, particularly longer inserts, are often missing, as in seen in **Figure 2E**. To investigate the structures of inserts, we used Alphafold models for 5,974 *C. albicans* proteins (Jumper et al. 2021; Varadi et al. 2022).

Alphafold models include a confidence score for each residue. The pLDDT (predicted local distance test) score measures the agreement of the prediction with an experimental model. The scores are on a scale of 0 to 100, where scores >90 are considered highly accurate, >70 correspond to a confident prediction with the correct backbone prediction. pLDDT scores <50 indicate a high likelihood of being unstructured, and can be used as a prediction of disorder (Tunyasuvunakool et al. 2021). It should be noted that Alphafold structures are of monomeric proteins and unstructured regions may become structured if the protein is part of a complex, as seen with Sth1 (below)

We extracted pLDDT scores for the residues in 3,013 of the 3,069 *C. albicans* inserts containing 21,811 residues from 1,007 Alphafold models. The average pLDDT score for all insert residues is 51.72. Only 8.9% of insert residues have pLDDT scores >90 and are predicted with high confidence (**Supplementary Table 3**). Just over one quarter of insert residues are predicted with confidence (pLDDT > 70; 5,689/21,811; 26.1%). Over half of insert residues (58.8%) have pLDDT scores <50 and have a high likelihood of being unstructured. Disordered proteins participate in many cellular signaling processes (Wright and Dyson 2015), functioning as hubs in protein interaction networks (Kim et al. 2008). Our identification of inserts in the majority of kinases additionally supports the idea of inserts playing a role in signaling.

How does the pLDDT score vary with insert length? The average pLDDT score for residues in inserts with 10 or fewer residues is 66.04 (**Supplementary Table 4**). This is the only range of insert length with an average > 50. Inserts of 11 or more residues are more likely to be unstructured. Inserts of 31 to 100 residues in length have the lowest average pLDDT scores. Inserts of these lengths may be too short to readily adopt stably folded structures and too long to be influenced by insert-flanking residues. Longer inserts, those over 100 residues in length are somewhat more likely to fold.

### Insertion sequences are flanked by structured regions

Into what structural context are insertion sequences inserted? The pLDDT scores of selected inserts of 10, 20 and 40 amino acids in length were extracted along with the scores of 20 flanking residues (**Figure 3**). For the shorter 10 residue inserts (top panel), most flanking residues have pLDDT scores >70, indicating they are confidently predicted and the flanking regions are folded. As inserts get longer, more flanking residues have pLDDT scores <50, suggesting they are disordered. All possible combinations of structured and unstructured inserts flanked by structured or unstructured residues can be found.

**Figure 3.**
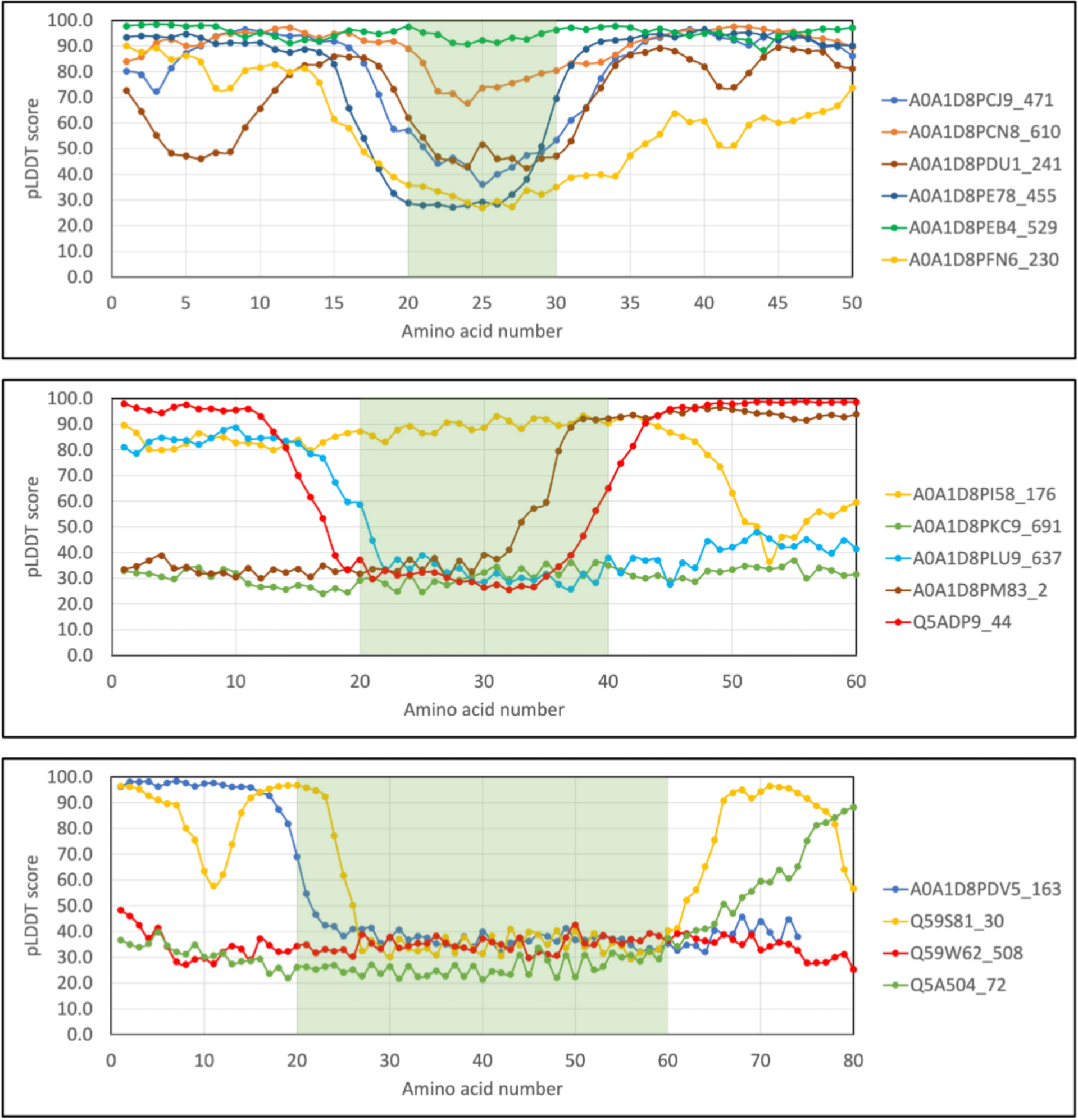
The pLDDT scores of inserts and flanking residues. The pLDDT scores of inserts with 10 (top), 20 (middle) and 40 (bottom) amino acids in length and 20 flanking amino acids N- and C-terminally were plotted. The colored rectangles highlight the extent of the insert. The legends indicate the UniProt accession of the protein followed by the number of the first amino acid plotted.

Inserts are most often unstructured sequences inserted into structured regions of the protein. The average pLDDT scores for the 59,420 N-terminal and 59,989 C-terminal insert-flanking residues are 77.49 and 77.38, respectively. Thus, both N- and C-terminal flanking residues are on average confidently predicted and likely folded, with pLDDT scores an average of ∼25 points higher than inserts themselves. Whereas 58.8% of insert residues have pLDDT scores <50, the N- and C-terminal fractions <50 are 18.2% and 18.1%, respectively (**Figure 4A**).

**Figure 4A.**
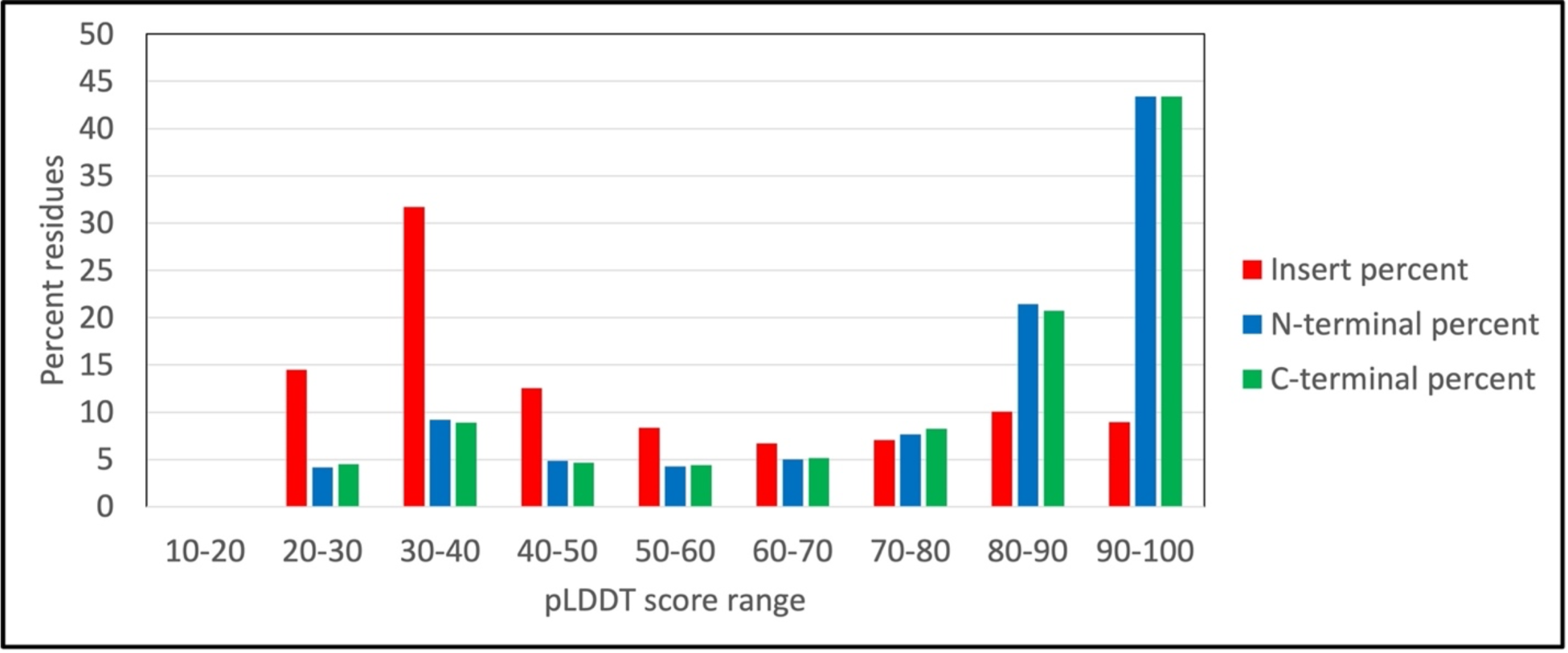
The distribution of pLDDT scores for residues in inserts or in flanking residues. The pLDDT scores insert residues and of 20 insert-flanking N- and C-terminal amino acids were extracted and binned by score range. The percentages of residues in each pLDDT score range are plotted for inserts (red), N-terminal flanks (blue) and C-terminal flanks (green).

### Insertion sequences are highly depleted of hydrophobic amino acids and highly enriched in polar amino acids

What is the amino acid composition of *C. albicans* inserts? Eukaryotic proteomes contain more disordered regions than prokaryotic proteomes (Basile et al. 2019). Eukaryotic proteins have more extended and more disordered linker regions (Basile et al. 2019). Changes in abundance of three amino acids, Ser, Pro and Ile were suggested to contribute to the increased disorder in eukaryotic proteins. We compared the amino acid compositions of 21,811 insert residues to the 704,020 residues of the 1,007 insert-containing proteins with Alphafold models (**Table 5**). Chi-square analysis was used to determine whether the observed number of each amino acid type was significantly different from an expected value. All pvalues are highly significant except for Lys. The most highly enriched amino acids in inserts are particularly Asn and Ser, but also Gln, Asp, Thr, Gly, Pro and His. The most highly depleted amino acids in inserts are particularly Leu, but also Ile, Phe, Val, Trp, Cys, Met, Tyr, Arg and Ala. Hydrophilic amino acids are significantly over-represented and hydrophobic amino acids are significantly under-represented in inserts, consistent with a surface location for most inserts. An important driving force in protein folding is the hydrophobic effect. The presence of hydrophobic amino acids in inserts on protein surfaces may interfere with the folding of newly synthesized copies of the protein by attempting to bury the inserted hydrophobic residues into the protein’s interior (Van Gils et al. 2022). In addition, hydrophobic inserts on the protein surfaces may promote inappropriate protein-protein interactions and are associated with diseases of protein aggregation (Van Gils et al. 2022). Inserts containing hydrophobic amino acids are therefore more likely to abolish or severely diminish protein function and will be preferentially eliminated or expressed at lower levels (Van Gils et al. 2022). Over-representation of Ser and Thr in inserts makes them targets for Ser/Thr protein kinases, again suggesting that inserts could play regulatory roles (Basile et al. 2019).

**Table 5.** Number of each amino acid in inserts and in the proteins they originate from. Amino acids in inserts and in the full-length proteins were tabulated. Chi-square analysis was used to compare the numbers for each amino acid type in inserts to the number in full-length proteins. Amino acids over-represented in inserts are highlighted.

| Amino Acid | Number in Inserts | Insert Percent | Number in Proteins | Protein Percent | p value |
| --- | --- | --- | --- | --- | --- |
| ALA | 936 | 4.3 | 38,639 | 5.5 | $3.16 \times 10^{-15}$ |
| ARG | 576 | 2.6 | 27,805 | 3.9 | $6.78 \times 10^{-24}$ |
| ASN | 2,297 | 10.5 | 43,504 | 6.2 | $6.22 \times 10^{-162}$ |
| ASP | 1,771 | 8.1 | 42,192 | 6.0 | $3.40 \times 10^{-41}$ |
| CYS | 63 | 0.3 | 7,576 | 1.1 | $2.42 \times 10^{-30}$ |
| GLN | 1,330 | 6.1 | 29,516 | 4.2 | $3.72 \times 10^{-46}$ |
| GLU | 1,677 | 7.7 | 45,913 | 6.5 | $1.32 \times 10^{-12}$ |
| GLY | 1,478 | 6.8 | 38,745 | 5.5 | $5.53 \times 10^{-17}$ |
| HIS | 518 | 2.4 | 14,359 | 2.0 | $3.71 \times 10^{-04}$ |
| ILE | 1,009 | 4.6 | 50,583 | 7.2 | $5.51 \times 10^{-50}$ |
| LEU | 1,193 | 5.5 | 66,938 | 9.5 | $8.88 \times 10^{-95}$ |
| LYS | 1,513 | 6.9 | 51,058 | 7.3 | $6.80 \times 10^{-02}$ |
| MET | 183 | 0.8 | 12,888 | 1.8 | $1.29 \times 10^{-28}$ |
| PHE | 533 | 2.4 | 31,599 | 4.5 | $1.16 \times 10^{-49}$ |
| PRO | 1,057 | 4.8 | 30,265 | 4.3 | $5.16 \times 10^{-05}$ |
| SER | 2,670 | 12.2 | 59,149 | 8.4 | $8.65 \times 10^{-96}$ |
| THR | 1,699 | 7.8 | 40,760 | 5.8 | $8.82 \times 10^{-38}$ |
| TRP | 47 | 0.2 | 7,276 | 1.0 | $6.91 \times 10^{-34}$ |
| TYR | 476 | 2.2 | 24,615 | 3.5 | $7.21 \times 10^{-27}$ |
| VAL | 785 | 3.6 | 40,640 | 5.8 | $2.02 \times 10^{-44}$ |
| HYDROPHOBES (ILV) | 2,987 | 13.7 | 158,161 | 22.5 | $3.64 \times 10^{-218}$ |
| AROMATICS (FHYW) | 1,574 | 7.2 | 77,849 | 11.1 | $2.04 \times 10^{-75}$ |
| ACIDIC (DE) | 3,448 | 15.8 | 88,105 | 12.5 | $1.94 \times 10^{-50}$ |
| BASIC (HRK) | 2,607 | 12.0 | 93,222 | 13.2 | $1.17 \times 10^{-08}$ |
| Total | 21,811 | 100.0 | 704,020 | 100.0 |  |

DisProt is a database of intrinsically disordered proteins (IDP) (Quaglia et al. 2022). Amino acids that are over-represented in IDP include Gln, Lys, Pro, Glu and Ser, while under-represented amino acids in IDP include Phe, Ala, Val, Ile, Leu as well as Arg and aromatic amino acids. There is considerable overlap in the sets of enriched or depleted amino acids between DisProt proteins and inserts. The most notable differences are the very high over-representation of Asn in inserts and the over-representation of Lys in IDP. The amino acid composition of inserts and their propensity to structural disorder is consistent with the very low pLDDT scores for inserts in Alphafold models. Inserts with sequences that strongly favor the adoption of a stable secondary structure may interfere with the folding of the remainder of the protein and be more strongly selected against.

### Insertion sequences can acquire functional significance

The *S. cerevisia*e Sth1 protein encodes the catalytic subunit of the RSC (Remodel the Structure of Chromatin) complex, which has vital roles in DNA repair and replication and in transcriptional regulation (Chen et al. 2020). Sth1 has 2 inserts, a 7-residue CDI (residues 222-227) and a 64-residue insert (residues 1,166-1,229). Sth1 residues 1,183-1,359 were shown to tether the Taf14 (Transcription initiation factor TFIID subunit 14) to the RSC complex (Kabani et al. 2005). Sth1 residues 1,183-1,240, which include much of the long Sth1 insert, were analyzed by nuclear magnetic resonance spectroscopy in complex with the C-terminal BET (pfam17035; <u>b</u>romodomain <u>e</u>xtra-<u>t</u>erminal) domain of Taf14 (residues 174-244) (**Figure 4B, Supplementary Figure 3C**) (Chen et al. 2020). The BET-binding motif is comprised of alternating hydrophobic and basic residues; in Sth1 residues 1,204-1,210 contain the motif. Sth1 forms a three-stranded ß-sheet with two strands from Taf14 and the third strand being the motif sequence. This motif has been conserved in all of the insert-containing sequences of the Saccharomycetaceae clade (**Supplementary Figure 3A**). The hydrophobic residues of the motif form extensive contacts with non-polar residues in Taf14 and the basic residues are oriented towards negatively charged surfaces on Taf14 (Chen et al. 2020). Disruption of the Sth1-Taf14 interaction alters Taf14 transcriptional regulation, with over 400 genes differentially regulated (Chen et al. 2020). Cells with the disruption are particularly sensitive to heat and osmotic stress (Chen et al. 2020).

**Figure 4B.**
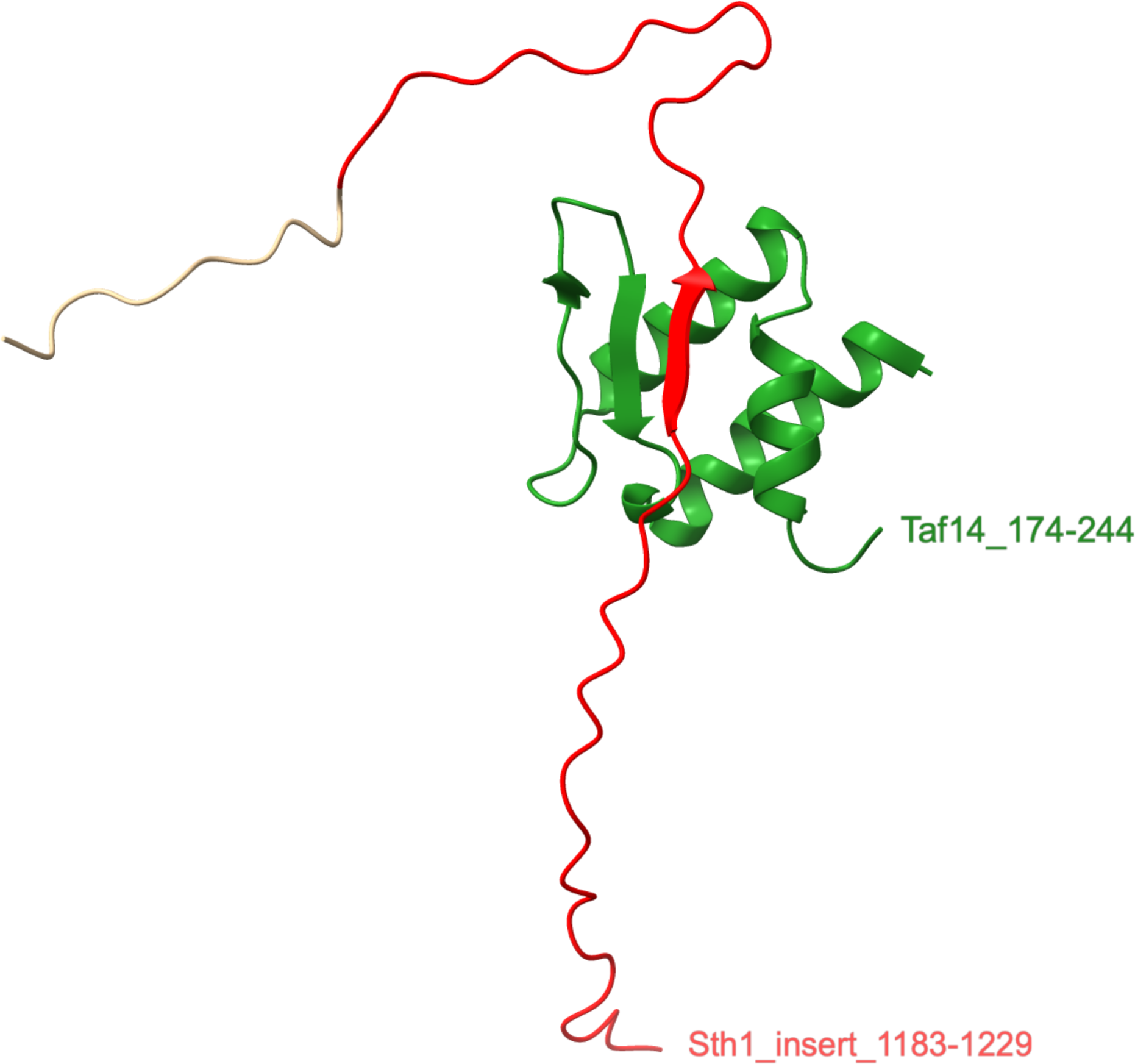
Structure of the Taf14-Sth1 complex. Chains 1.1_A (Taf 14, green) and 1.1_B (Sth1, red and tan) were selected from the ensemble of chains in PDB: 6LQZ. The Sth1 insert residues 1,183-1,229 are shown in red.

Sth1 also contains a second insert, residues 222-227 involved in protein-protein interactions with RSC subunits, RSC2 and RSC58. The insert extends as a loop into a pocket formed by the RSC2 and RSC58 subunits (**Supplementary Figure 3B**). The functional significance of this interaction, which has a buried surface area of 415 Å^2^ is unknown. Thus, newly acquired inserts can modify or expand protein function.

### Conclusions

Our study revealed a large number of inserts in protein-coding regions in Ascomycete fungi. The total number of inserts in each clade correlates with the rate of evolutionary change, suggesting that similar mechanisms of insert generation are involved in all four clades. CDIs numbers are greater in budding yeasts than in filamentous fungi, suggesting differences in insert retention rather than insert generation may be responsible.

How are inserts and deletions generated? We have investigated 3n indels ranging from three to several hundred base pairs in protein-coding regions. The diversity of indels is consistent with a variety of mutagenic events, including double-strand breaks, which can be repaired by homologous recombination and non-homologous end joining and give rise to insertions or deletions of variable length (Hanscom and McVey 2020). Which mechanisms are employed to repair DNA damage will necessarily be dependent on the presence of repair proteins in the genome. Determining which mechanisms are most responsible for generating inserts will require further investigation, but will be hampered by the fact that the DNA surrounding the original mutagenic event will have continued to undergo mutations for millions of years.

Indels in coding regions are expected to be under strong selective pressure. The indels we studied were benign or at least insufficiently deleterious to protein function to allow strain survival and indel inheritance. Indels are largely located on protein surfaces, suggesting that indels occurring within protein cores are too disruptive to protein structure or function. Over time, retained indels have been subject to additional mutation events, providing opportunities for proteins to develop new properties or even new functions.

One Sth1 insert (1,166-1,229) has acquired a central role in that protein’s function. It is interesting to note that all the residues of this insert have pLDDT scores <50 and are predicted to be unstructured. Yet, in PDB: 6LQZ, protein-protein interactions result in a portion of the insert adopting a ß-strand conformation. Although we found most inserts are predicted to be unstructured, complex formation may reveal new conformations. Thus, it remains to be determined how many inserts are functionally important. The challenge will be to identify which retained indels have evolved functional roles.

## METHODS

All proteomes were downloaded from NCBI RefSeq in FASTA format (O’Leary et al. 2016). Only one proteome was used per species. Database accessions and the number of proteins per proteome are available in **Supplementary Excel file 1**. A web server called Bagheera was used with *C. albicans* as the reference species to determine the most probable CUG codon translation for the *S. spartinae* genome, GCF_019049425.1_ASM1904942v1_genomic.fna (**Supplementary Discussion)** (Mühlhausen and Kollmar 2014; Villarreal et al. 2021).

The 6,030 proteins of *C. albicans SC5314* served as query proteins in blastp searches of the 81 chosen proteomes. A single best hit with an evalue less than 9.9 e^-30^ was retained (**Archive_fasta)**. MSAs were generated using MAFFT v7.475 with default parameter except the leavegappyregion option was employed (**Archive_MSAs)** (Katoh and Standley 2016). MAFFT with these parameters was found to produce MSAs with fewer gaps than other algorithms. Phylogenetic inferences using maximum likelihood were performed with IQ-TREE 2 (Minh et al. 2020).

1. *C. albicans* insert-containing proteins were analyzed for Pfam (Pfam 35.0) domains using the Conserved Domain Database with an evalue cutoff of 9.99e-06 (Lu et al. 2020; Mistry et al. 2021). Orthology analysis used the KEGG KofamKOALA web server (KEGG release 101.0) (Aramaki et al. 2020). Some proteins are assigned to more than one orthologous group or functional classification; in determining the numbers of proteins assigned to each classification, a single assignment for each protein was used. To investigate the effects of inserts on protein-protein interactions, we used STRING (version 11.5) (Szklarczyk et al. 2021). Complex Portal was used to investigate the effects of inserts on membership in macromolecular complexes in *S. cerevisiae*, the only organism in our study currently included in the database (Meldal et al. 2021). The Alphafold database contains 5,974 *C. albicans* protein models (May, 2022); proteins longer than 2,700 amino acids are not included. Models were downloaded and the pLDDT scores extracted from the mmCIF files (Varadi et al. 2022). Protein structure images were generated in Chimera (v1.25) (Pettersen et al. 2021).

## Supporting information

Supplementary discussion

fasta protein sequences

MAFFT_MSAs

## Acknowledgements

This work was supported by NIH grants R01AI141794 and R21AI69276 awarded to PU. B.D.L. would like to thank Bart Hazes for his invaluable help with python programming.

## Author contributions

B.D.L. and P.U. conceived the study and wrote the manuscript. B.D.L. executed the data acquisition and analysis.

