## Supplementary tables and figures for "Coding Sequence Insertions in Fungal Genomes are Intrinsically Disordered and can Impart Functionally-Important Properties on the Host Protein"

| CTG strains | No. CDIs | Saccharomycetaceae strains | No. CDIs | Aspergillaceae strains | No. CDIs | Herpotrichiellaceae strains | No. CDIs |
|---|---|---|---|---|---|---|---|
| Candida albicans | 1366 | Candida glabrata | 1466 | Aspergillus aculeatus | 226 | Capronia coronata | 431 |
| Candida auris | 1332 | Eremothecium cymbalariae | 1319 | Aspergillus clavatus | 221 | Capronia epimyces | 439 |
| Candida dubliniensis | 1364 | Eremothecium gossypii | 1325 | Aspergillus fischeri | 220 | Cladophialophora bantiana | 495 |
| Candida duobushaemulonis | 1295 | Kazachstania africana | 1472 | Aspergillus flavus | 235 | Cladophialophora carrionii | 493 |
| Candida haemuloni | 1327 | Kazachstania naganishii | 1468 | Aspergillus fumigatus | 219 | Cladophialophora immunda | 480 |
| Candida orthopsilosis | 1310 | Kluyveromyces lactis | 1362 | Aspergillus nidulans | 237 | Cladophialophora psammophila | 494 |
| Candida pseudohaemulonii | 1316 | Kluyveromyces marxianus | 1358 | Aspergillus niger | 207 | Cladophialophora yegresii | 483 |
| Candida tropicalis | 1296 | Lachancea lanzarotensis | 1349 | Aspergillus nomiae | 232 | Exophiala aquamarina | 427 |
| Clavispora lusitaniae | 1266 | Lachancea thermotolerans | 1350 | Aspergillus oryzae | 223 | Exophiala dermatitidis | 424 |
| Debaryomyces fabryi | 1273 | Naumovozyma castellii | 1498 | Aspergillus ruber | 218 | Exophiala mesophila | 413 |
| Debaryomyces hansenii | 1309 | Naumovozyma dairenensis | 1512 | Aspergillus sydowii | 247 | Exophiala oligosperma | 427 |
| Hyphopichia burtonii | 1336 | Saccharomyces cerevisiae | 1514 | Aspergillus terreus | 225 | Exophiala spinifera | 433 |
| Lodderomyces elongisporus | 1261 | Saccharomyces eubayanus | 1506 | Penicilliopsis zonata | 224 | Exophiala xenobiotica | 425 |
| Metschnikowia bicuspidata | 1299 | Saccharomyces paradoxus | 1503 | Penicillium arizonense | 264 | Fonsecaea erecta | 490 |
| Meyerozyma guilliermondii | 1225 | Tetrapisispora blattae | 1449 | Penicillium digitatum | 290 | Fonsecaea monophora | 489 |
| Scheffersomyces spartinae | 1228 | Tetrapisispora phaffii | 1482 | Penicillium expansum | 289 | Fonsecaea multimorphosa | 488 |
| Scheffersomyces stipitis | 1359 | Torulaspora delbrueckii | 1453 | Penicillium griseofulvum | 289 | Fonsecaea nubica | 490 |
| Spathaspora passalidarum | 1274 | Vanderwaltozyma polyspora | 1421 | Penicillium roqueforti | 289 | Fonsecaea pedrosoi | 490 |
| Suhomyces tanzawaensis | 1336 | Zygosaccharomyces rouxii | 1441 | Penicillium rubens | 289 | Phialophora attinorum | 347 |
| Yamadazyma tenuis | 1221 | Zygotorulaspora mrakii | 1468 | Penicillium solitum | 293 | Rhinocladiella mackenziei | 442 |
| CDIs shared by all strains | 640 (39.3%) | CDIs shared by all strains | 881 (50.5%) | CDIs shared by all strains | 199 (55.3%) | CDIs shared by all strains | 291 (52.4%) |

**Supplementary Table 1. Number of CDIs retained by each strain of the four taxonomic families**. CDIs were counted as retained if at least one amino acid was present in the length of the insertion event. The highest and lowest numbers of retained CDIs are highlighted for each clade in green and red, respectively. The number of CDIs retained by all 20 strains of each clade is also indicated. The number of CDIs retained as a fraction of the total number of CDIs in the clade is presented as a percentage.

| Strain | *C. albicans* | *S. cerevisiae* | *A. oryzae* | *C. coronata* |
|---|---|---|---|---|
| inserts total | 3,069 | 3,193 | 1,261 | 1,782 |
| insert length 1-5 | 2,056 | 2,154 | 835 | 1,182 |
| insert length 6-10 | 449 | 463 | 223 | 306 |
| insert length 11-20 | 314 | 314 | 143 | 198 |
| insert length 21-50 | 202 | 219 | 54 | 83 |
| insert length >50 | 48 | 43 | 6 | 13 |
| insert length >100 | 8 | 9 | 1 | 4 |
| no. of MSAs | 1,015 | 1,064 | 623 | 736 |
| inserts per MSA | 3.0 | 3.0 | 2.0 | 2.4 |
| % MSAs with inserts | 66.3 | 69.5 | 40.7 | 48.0 |

**Supplementary Table 2. Number of inserts present in one strain from each clade.** Inserts in each of the four strains were identified and grouped by length. The total number of MSAs analyzed is 1,532 and the number of MSAs in which all inserts are found is indicated. Only MSAs with inserts are included in the calculation of inserts per MSA.

| pLDDT score | No. of residues | Percent |
|---|---:|---:|
| 10-20 | 17 | 0.1 |
| 20-30 | 3,164 | 14.5 |
| 30-40 | 6,918 | 31.7 |
| 40-50 | 2,734 | 12.5 |
| 50-60 | 1,823 | 8.4 |
| 60-70 | 1,466 | 6.7 |
| 70-80 | 1,541 | 7.1 |
| 80-90 | 2,196 | 10.1 |
| 90-100 | 1,952 | 8.9 |
| **Total** | **21,811** | **100.0** |

**Supplementary Table 3. Number of insert residues in each pLDDT score range**. The pLDDT scores for 21,811 residues from 3,013 inserts were extracted and number of residues in each score range tabulated.

| Insert length | No. of inserts | Average pLDDT score | No. of residues |
|---|---:|---:|---:|
| 1-10 | 2,458 | 66.04 | 7,591 |
| 11-20 | 308 | 48.17 | 4,477 |
| 21-30 | 112 | 44.80 | 2,778 |
| 31-40 | 60 | 39.95 | 2,085 |
| 41-50 | 27 | 39.76 | 1,223 |
| 51-100 | 40 | 40.25 | 2,591 |
| 101-150 | 8 | 47.40 | 1,066 |
| **Total** | **3,013** | **51.72** | **21,811** |

**Supplementary Table 4. Average pLDDT scores by insert length**. The pLDDT scores for 21,811 residues from 3,013 inserts were extracted and average pLDDT score for each insert range tabulated.

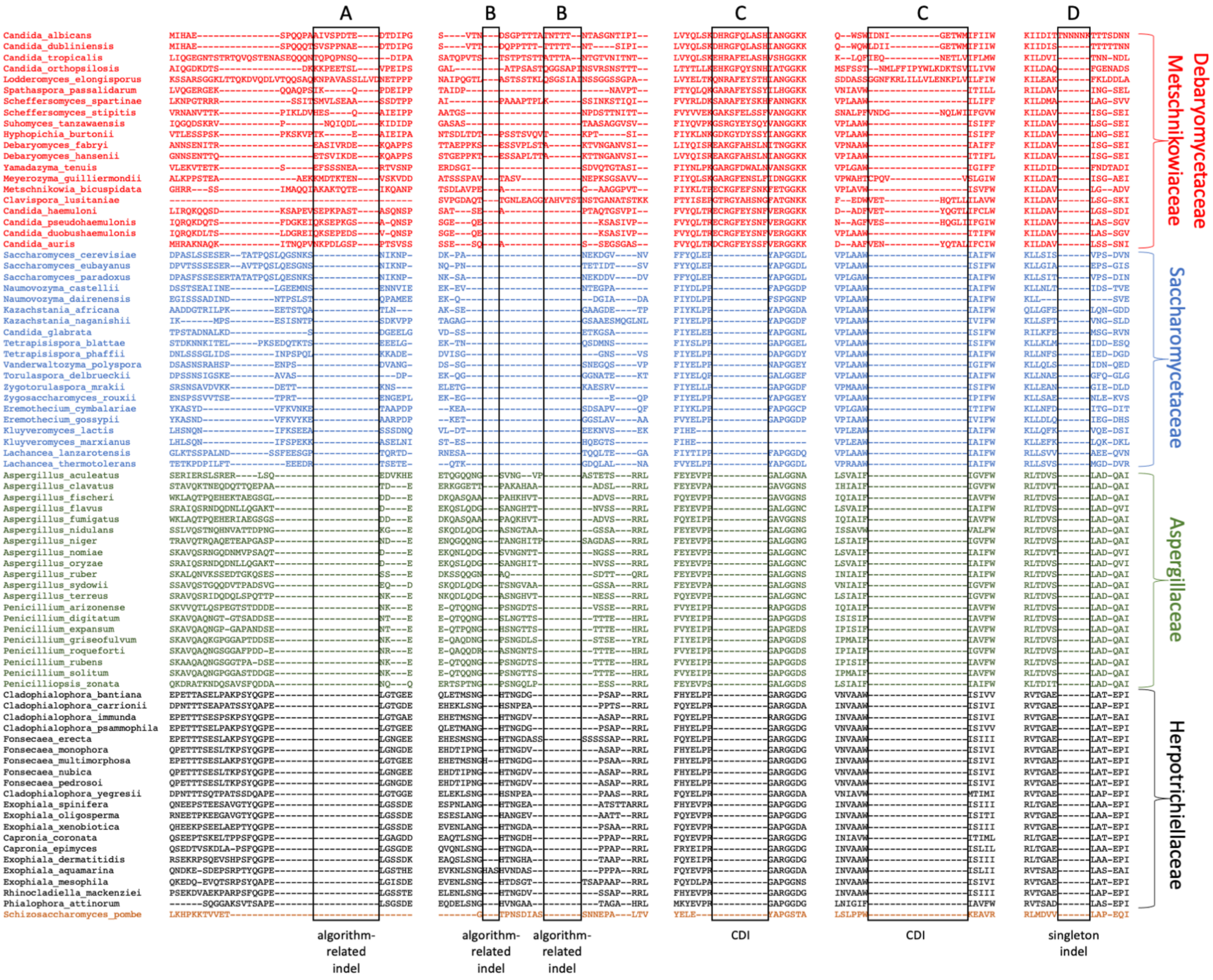
A
B
B
C
C
D
Debaryomycetaceae
Metschnikowiaceae
Saccharomycetaceae
Aspergillaceae
Herpotrichiellaceae
algorithm-related indel
algorithm-related indel
algorithm-related indel
CDI
CDI
singleton indel

**Supplementary Figure 1A. Examples of inserts.** Five partial alignments are shown. (**A,B**) Algorithm-related inserts are at least partly generated from matrix scoring and/or gap insertion parameters of the algorithm. (**C**) Clade-defining inserts (CDIs) have inserts restricted to a single clade. The CDI must have at least one position with 5 or more aligned amino acids, but otherwise may contain gaps. CDIs must have well-aligned flanking positions with 59 or more residues on both sides of the indel. (**D**) Singleton inserts are found in only one sequence.

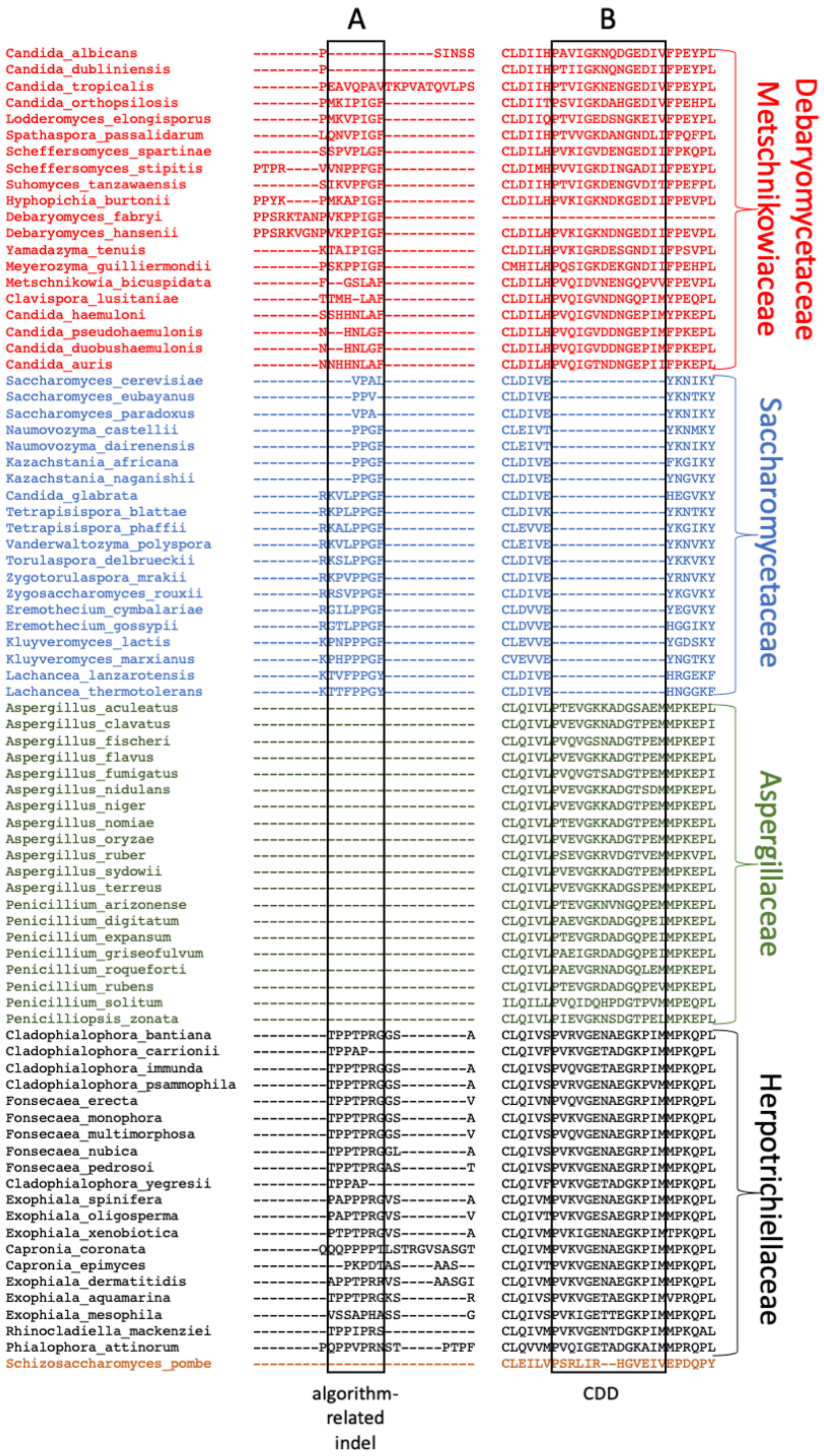
A
B
Debaryomycetaceae
Metschnikowiaceae
Saccharomycetaceae
Aspergillaceae
Herpotrichiellaceae
algorithm-related indel
CDD

**Supplementary Figure 1B. Examples of deletions.** Two partial alignments are shown. (**A**) An algorithm-related deletion in Aspergillaceae likely arose from the insertion of gaps into a poorly aligning region of the MSA. (**B**) Clade-defining deletions (CDDs) have deletions restricted to a single clade, here in Saccharomycetaceae. The CDD must have at least 5 amino acids in each of the three other clades, but otherwise may contain gaps. CDDs must have flanking positions with 40 or more residues on both sides.

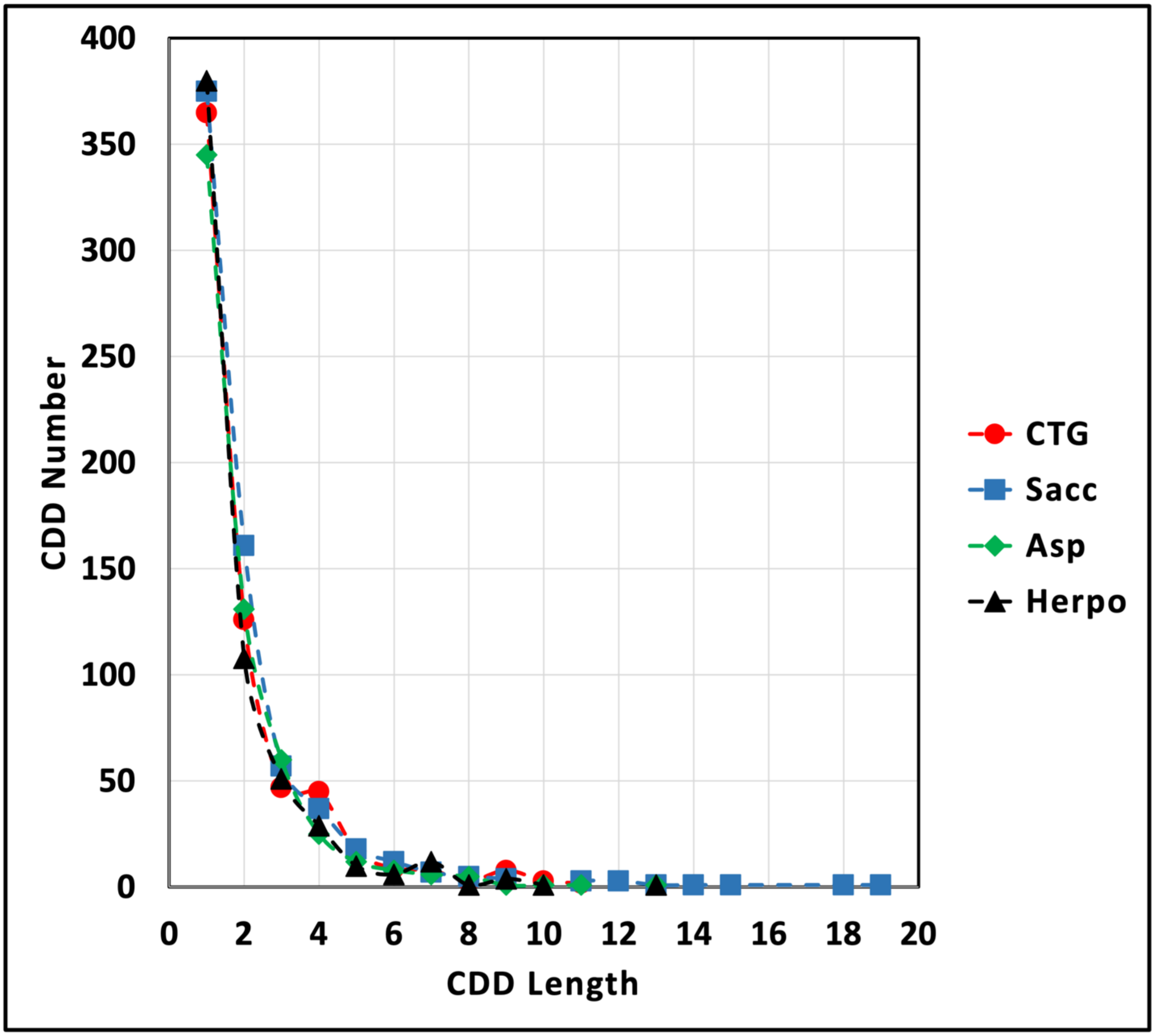


**Supplementary Figure 2A. Length distribution of CDDs**. The numbers of CDDs were plotted for each clade. Red circles, CTG clade; blue squares, Saccharomycetaceae clade; green diamonds, Aspergillaceae clade; black triangles, Herpotrichiellaceae clade.

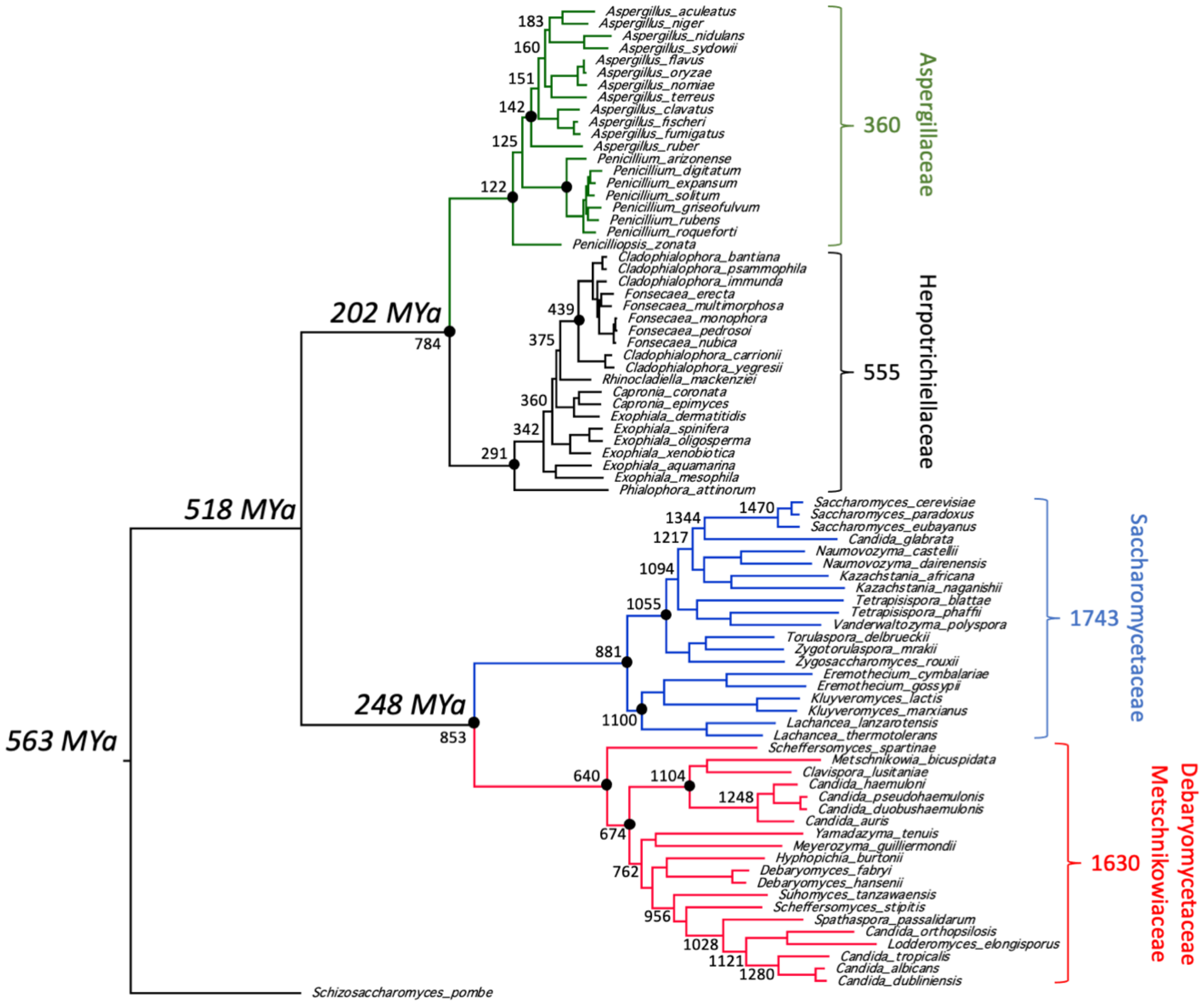


**Supplementary Figure 2B. CDI retention within clades**. The evolutionary times associated with selected nodes were derived from a recent comprehensive investigation of Ascomycota phylogeny (Shen et al. 2020). Selected nodes with 100% bootstrap support are indicated with black circles. The total numbers of CDIs for each clade are indicated by the taxonomic labels. The numbers associated with selected nodes are the numbers of shared CDIs by all descendants from that node. The numbers of inserts that are shared by the CTG and Saccharomycetaceae clades and by the Aspergillaceae and Herpotrichiellaceae clades are also indicated.

| | |
|---|---|
| NP_012140_Saccharomyces_cerevisiae/1203-1210 | KLTVKIKL |
| XP_018221496_Saccharomyces_eubayanus/1203-1210 | KLTVKIKL |
| XP_033766929_Saccharomyces_paradoxus/1202-1209 | KLTVKIKL |
| XP_003675109_Naumovozyma_castellii/1170-1177 | KLKIKLTM |
| XP_003668661_Naumovozyma_dairenensis/1196-1203 | KLKIKLTL |
| XP_003959368_Kazachstania_africana/1172-1179 | KFKITLKL |
| XP_022464686_Kazachstania_naganishii/1200-1207 | RLRVKLTL |
| XP_446743_Candida_glabrata/1198-1205 | KLKIKLKL |
| XP_004181716_Tetrapisispora_blattae/1231-1238 | KFKVKLTL |
| XP_003686053_Tetrapisispora_phaffii/1243-1250 | SLKVKISL |
| XP_001646124_Vanderwaltozyma_polyspora/1197-1204 | KLKIKLKL |
| XP_003680552_Torulaspora_delbrueckii/1162-1169 | KLKIKLTL |
| XP_037144276_Zygotorulaspora_mrakii/1188-1195 | KLKIRLKL |
| XP_002494432_Zygosaccharomyces_rouxii/1197-1204 | KLKIKLNL |
| XP_003646975_Eremothecium_cymbalariae/1171-1178 | KLKVKISL |
| NP_985231_Eremothecium_gossypii/1150-1157 | KLKVKINL |
| XP_455284_Kluyveromyces_lactis/1215-1222 | KIKLKLNM |
| XP_022677605_Kluyveromyces_marxianus/1198-1205 | KIKLKLNM |
| XP_022628368_Lachancea_lanzarotensis/1151-1158 | RLTVKLSL |

**Supplementary Figure 3A. Taf14-binding motif in Saccharomycetaceae clade inserts**. Basic residues are highlighted in red; hydrophobic residues are highlighted in cyan. NP_012140 is the accession code for Sth1.

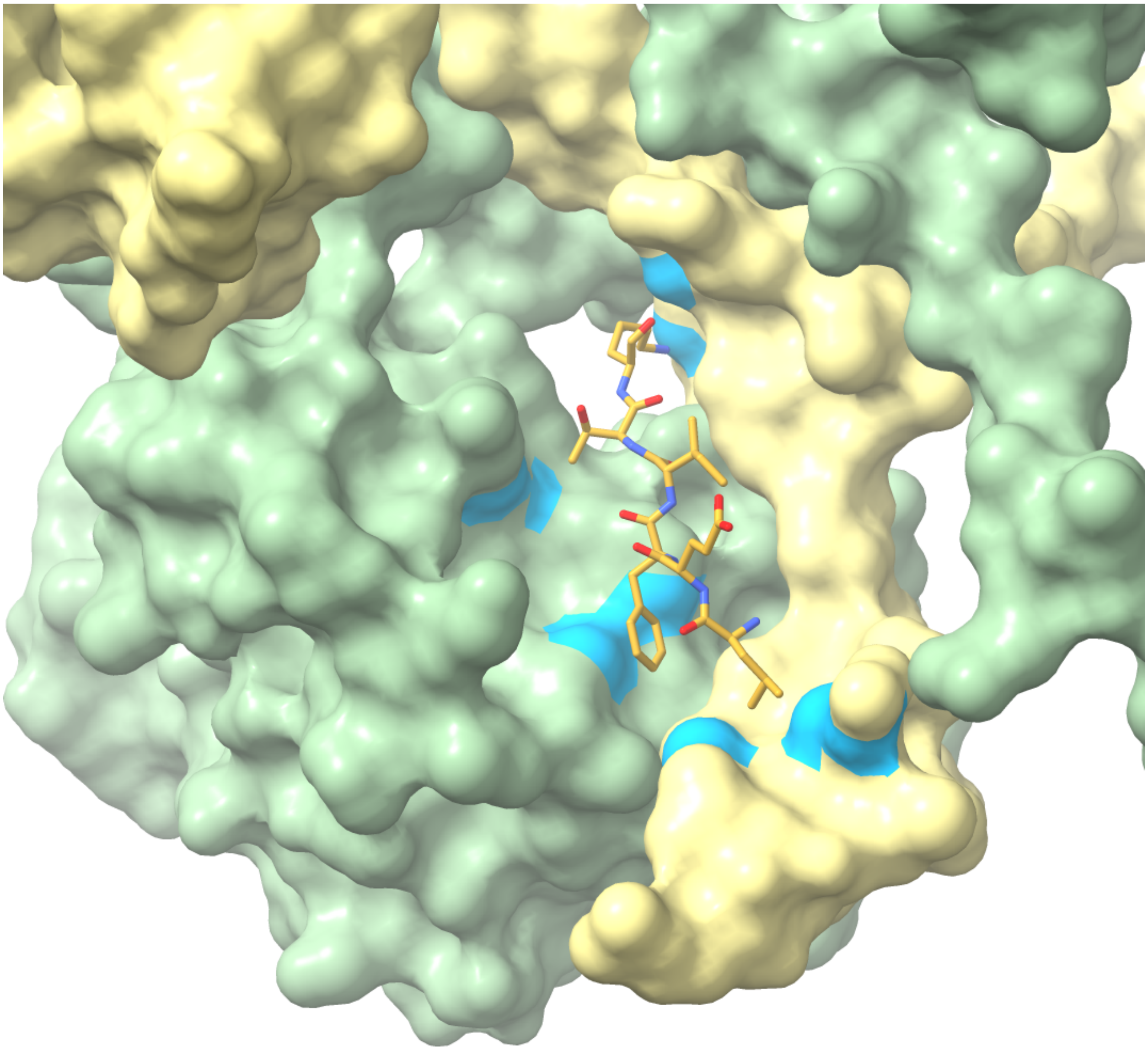

**Supplementary Figure 3B**. **Sth1 insert 222-227 is located in a pocket formed by RSC2 and RSC58.** The structure accession is PDB:6K15 (Chen et al. 2020)**.** Sth1 residues 222-227 are depicted in gold as a stick model. RSC2 and RSC58 are depicted in surface representation (khaki and green, respectively). Surfaces with 4 Å of residues 222-227 are colored blue.

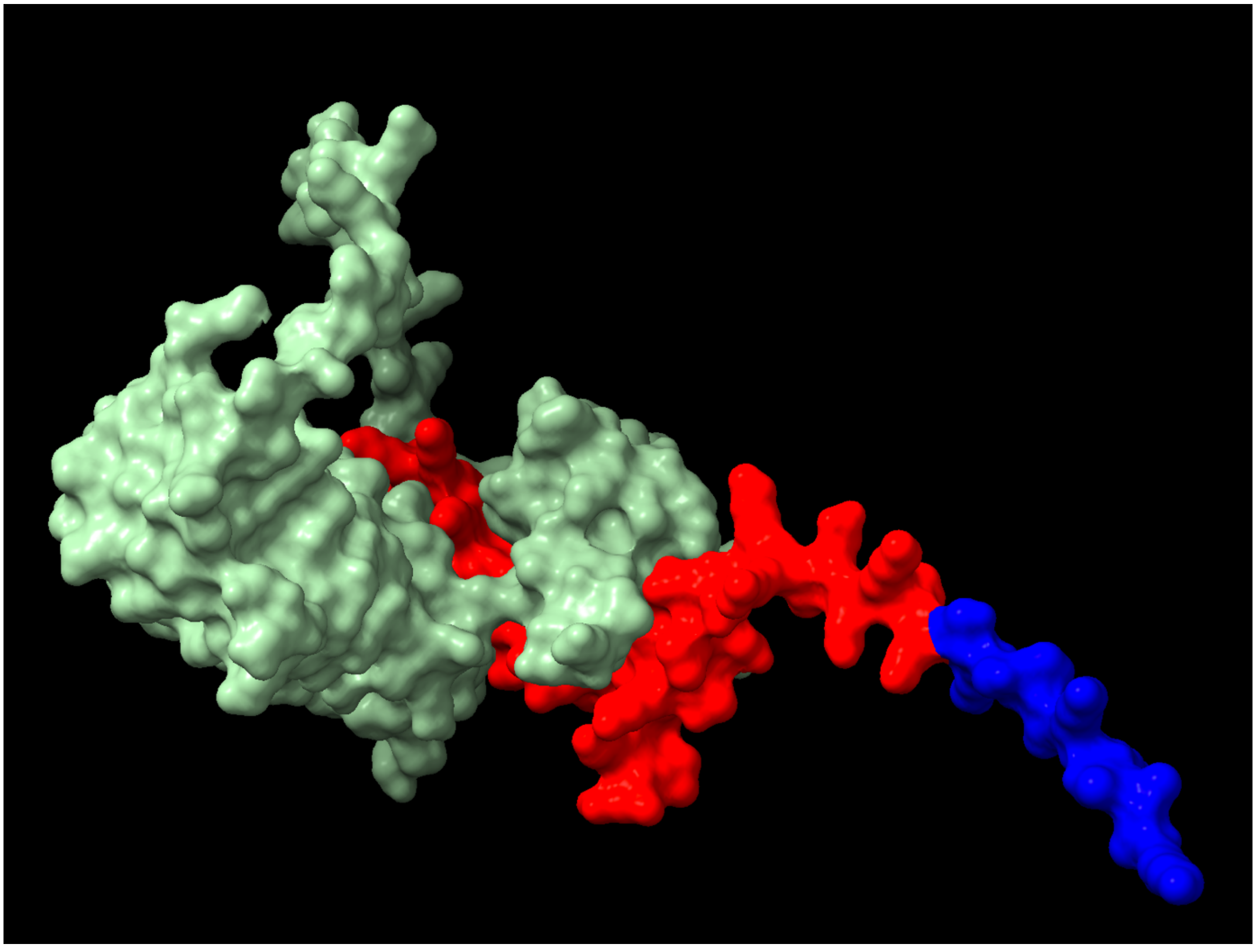

**Supplementary Figure 3C. Interaction of the Sth1 Insert Residues 1,183-1,229 with Taf14.**

Chains 1.1_A (Taf 14, green) and 1.1_B (Sth1, red and blue) were selected from the ensemble of chains in PDB: 6LQZ for display in a surface representation. The Sth1 insert residues 1,183-1,229 are shown in red.
