## Supplementary discussion for "Coding Sequence Insertions in Fungal Genomes are Intrinsically Disordered and can Impart Functionally-Important Properties on the Host Protein"

### *Scheffersomyces spartinae*

The genome sequence of *Scheffersomyces spartinae ARV011* was a new NCBI RefSeq genome deposited in June, 2021 (Villarreal et al., 2021). The organism was isolated from marine water and sediment in Massachusetts, USA and is yet poorly characterized. NCBI taxonomy initially indicated that this yeast utilizes the standard genetic code, rather than the alternative yeast genetic code (AYC), which translates CUG codons to serine rather than leucine (Schoch et al., 2020). Bagheera is a web server that determines the most probable CUG codon translation of genome sequences (Mühlhausen and Kollmar, 2014). Homologs of a selected set of cytoskeletal and motor proteins belonging to 79 *Saccharomycetes* yeast species serve as reference data to identify CUG codons and amino acid conservation. The reference database contains 8,244 known CUG codons, which are used to predict the most probable codon usage for every CUG codon found. For *S. spartinae*, homologs were identified for 34 of 38 reference proteins, 29 of which contained 159 CTG codons. Of these, 134 CTG codons were unambiguously aligned to reference data. The server predicts 54 codons are translated using the AYC, 12 are translated using the standard genetic code and 78 were indiscriminative. This is a strong signal that *S. spartinae* uses the AYC. For comparison, the authors analyzed the genome of *Candida maltosa Xu316*; 71 CTG codons were found in 20 proteins and 35 of these are conserved CTG positions translated with the AYC (Mühlhausen and Kollmar, 2014). The server did not identify the *S. spartinae* Ser-$tRNA_{CAG}$ involved in the alternative translation events, but this is not unusual, as introns may be present in the gene. *S. spartinae* uses the AYC and the NCBI taxonomy entry has been revised. Confirmation of this genetic code awaits biochemical investigation. Phylogenetically, *S. spartinae* belongs to the CTG clade.

**CDI retention and generation**

CDIs likely result from genetic changes that first occurred in a common ancestor of the group and were vertically inherited (Sharma and Gupta, 2019). In some cases, the CDI is lost and absent from the now extant group members. For the CTG clade, a total of 1,630 CDIs were identified (**Table 2**). The simplest explanation is that these insertions were generated after the evolutionary split of the CTG and Saccharomycetaceae clades, estimated to have occurred ~248 MYa (**Figure 1**). The last common ancestor of the 20 CTG strains used here would have had 1,630 CDIs. Since that time, speciation has occurred and some strains will have lost certain CDIs. We determined the number of CDIs present in each modern CTG strain. Although the length and amino acid composition of a CDI may change since its creation, a CDI was counted as being retained if at least one amino acid was present over the length of the insertion event.

For the 1,630 CDIs in the CTG strains, the numbers of retained CDIs range from 1,221 (74.9%) in *Yamadazyma tenuis* to 1,366 (83.8%) in *C. albicans* (**Supplementary Table 1**). For *C. albicans*, this means that 264 CDIs have been lost since the last common ancestor, which can be estimated to have been at ~126 MYa, or the time *S. spartinae* diverged from the other CTG strains. This corresponds to the loss of 2.1 CDIs/MY. For *Y. tenuis*, 409 CDIs have been lost (3.2 CDIs/MY).

In the Saccharomycetaceae clade, *Lachancea thermotolerans* and *S. cerevisiae* are estimated to have last had a common ancestor ~123.7 MYa (Shen et al., 2020) and this ancestor had 1,743 CDIs (**Supplementary Table 1**). For *S. cerevisiae*, 229 CDIs have been lost over that time (1.9 CDIs/MY). For *L. thermotolerans*, 393 CDIs have been lost (3.2 CDIs/MY). In the Aspergillaceae clade, *Penicillium zonata* and *A. oryzae* are estimated to have last had a common ancestor ~74.4 MYa and this ancestor had 360 CDIs (**Supplementary Table 1**). For *A. oryzae* and *P. zonata*, 137 and 136 CDIs have been lost, respectively (1.8 CDIs/MY). In the Herpotrichiellaceae clade, *Phialophora attinorum* and *Capronia bantiana* are estimated to have had a last common ancestor ~85.8 MYa, which had 555 CDIs (**Supplementary Table 1**). For *C. bantiana*, 60 CDIs have been lost (0.7 CDIs/MY). For *P. attinorum*, 208 CDIs have been lost (2.4 CDIs/MY). The rate of CDI loss differs between strains but the rates of loss differ as much between strains in the same clade as between clades.

We determined and recorded the numbers of shared CDIs on the phylogenetic tree (**Supplementary Figure 2B**). The numbers associated with selected nodes are the numbers of shared CDIs by all descendants from that node. For example, *C. albicans*, *C. dubliniensis* and *C. tropicalis* share 1,280 CDIs and 640 CDIs are shared by all CTG strains. We also determined inserts that are shared by the CTG and Saccharomycetaceae clades and by the Aspergillaceae and Herpotrichiellaceae clades. For example, these inserts must be found in at least five CTG and five Saccharomycetaceae strains but absent from all 41 other strains in the other two clades. The two budding yeast clades share 853 inserts, while the two filamentous fungal clades share 784. The most parsimonious explanation for these shared inserts is that the insertion event occurred after the last common ancestor of the four clades (518 MYa), but before the split between the clades (248 MYa for the budding yeast and 202 MYa for the filamentous fungi). From the time of the last common ancestor for all 4 clades (518 MYa) to the time that the two budding yeast clades diverged (248 MYa), 853 inserts were generated over 270 MY, or

3.2 inserts per MY; for filamentous fungi, 784 inserts were generated over 316 MY, or 2.5 inserts per MY. It is interesting to note that the rate of CDI loss (0.7-3.2 CDIs/MY) is comparable to the rate of insert generation (2.5-3.2 inserts/MY).
